# Delay of innate immune responses following influenza B virus infection affects the development of a robust antibody response in ferrets

**DOI:** 10.1101/2024.02.19.580935

**Authors:** Thomas Rowe, Ashley Fletcher, Robert A. Richardson, Melissa Lange, Giuseppe A. Sautto, Greg A. Kirchenbaum, Yasuko Hatta, Gabriela Jasso, David E. Wentworth, Ted M. Ross

## Abstract

Due to its natural influenza susceptibility, clinical signs, transmission, and similar sialic acid residue distribution, the ferret is the primary animal model for human influenza research. Antibodies generated following infection of ferrets with human influenza viruses are used in surveillance to detect antigenic drift and cross-reactivity with vaccine viruses and circulating strains. Inoculation of ferrets, with over 1500 human clinical influenza isolates (1998-2019) resulted in lower antibody responses (HI<1:160) to 86% (387/448) influenza B viruses (IBV) compared to 2.7% (30/1094) influenza A viruses (IAV). In this directed analysis, we show that the immune responses in ferrets inoculated with IBV (B/Victoria or B/Yamagata lineage) were delayed and reduced compared to IAV (A/H1pdm09 or A/H3N2). Analysis of innate gene expression in the ferret upper respiratory tract and peripheral blood indicated that IAV generated a strong inflammatory response, including an early activation of the interferon (IFN) response; whereas IBV elicited a delayed and reduced response. Serum levels of cytokines, chemokines, and IFNs were all much higher following IAV-infection than IBV-infection in ferrets. Pro-Inflammatory (MCP-1, MIP-1B, TNFA), IFN (IFNB, IFNG), TH1/TH2 (IP-10, IL-2), and T-effector (IL-12p40) proteins were significantly higher in sera of IAV-infected than IBV-infected ferrets over twenty-eight days following challenge. Serum levels of Type-I/II/III IFNs were detected following IAV-infection throughout the 28-day period and Type-III IFN was only detected by day 28 for IBV. An early increase in IFN-lambda levels corresponded to gene expression following IAV-infection. Reduced innate immune responses detected following IBV-infection reflected the subsequent delayed and reduced serum antibodies. Differences in serum antibody responses by IBV were not observed in antibody secreting cells in the spleen or peripheral blood. These findings help in understanding the antibody responses in humans following IBV vaccination or infection and consideration of potential addition of innate immunomodulators to overcome low responses.

**AUTHOR SUMMARY:** The ferret is the primary animal model for human influenza research. Using a ferret model, we studied the differences in both innate and adaptive immune responses following infection with influenza A and B viruses. Antibodies generated following infection of ferrets is used for surveillance assays to detect antigenic drift and cross-reactivity with vaccine viruses and circulating influenza strains. Influenza A virus (IAV) infection of ferrets to generate these reagents resulted in a strong antibody response, but Influenza B virus (IBV) infection generated weak antibody responses. In this study using influenza-infected ferrets, we found that IAV resulted in an early activation of the interferon and pro-inflammatory response, whereas IBV showed a delay and reduction in these responses. Serum levels of interferons and other cytokines or chemokines were much higher in ferrets following IAV infection. These reduced innate responses were reflected the subsequent delayed and reduced antibody responses to IBV in the sera. These findings may help in understanding low antibody responses in humans following influenza B vaccination and infection and may warrant the use of innate immunomodulators to overcome these weak responses.

## INTRODUCTION

Influenza A and B viruses circulate globally and represent a major public health concern. Prevention from infection or severe illness relies on a robust adaptive immune response. Antibodies following infection or vaccination are used in surveillance assays to detect antigenic drift and make recommendations for vaccine strain selection. This study involves the use of ferrets to better understand why influenza B viruses generate weak immune responses following infection. Ferrets are an ideal model to explore immune responses to influenza virus because they are naturally susceptible to influenza including human influenza strains without the need for prior adaptation (1, 2). Moreover, ferrets exhibit similar lung physiology and distribution of α2,3- and α2,6-linked sialic acids (3, 4); which are the receptors for influenza binding to cells. Finally, the frequency of antibody light chain isotypes in ferrets closely resembles that seen in humans (5) and further supports the use of ferrets for studying the antibody response elicited by influenza virus infection or vaccination.

Influenza A virus (IAV) and influenza B virus (IBV)-infected ferrets are used to generate antisera for influenza surveillance and to develop and improve influenza vaccines. Over a twenty-year period from 1998 to 2019, more than 1500 antisera raised to IAV and IBV have been generated in ferrets. In serology assays, such as the hemagglutination inhibition (HI) assay, reference antisera requires a minimal titer of at least 1:160 to adequately measure antigenic variation among viruses. For IBV, 86.4% of viruses (387/448) inoculated into ferrets required a booster dose to reach the HI titer of 1:160 compared to only 2.7% (30/1094) for IAV.

Low antibody responses in humans infected with IBVs have also been observed. A cohort study conducted in Hong Kong from 2009-2014 to assess the antibody responses to B-Vic and B-Yam infections showed that <50% of PCR-confirmed individuals had detectable neutralizing antibody titers. Additionally, all B-Vic and B-Yam infected individuals with antibody responses showed low titers (<160) (6). The effectiveness of IBV vaccine was also shown to be low, especially in children. In a study of the 2014/2015 influenza vaccine, which overall had 57% vaccine efficacy, it was shown that <20% of older children had a 4-fold rise in HI titer to influenza B following vaccination (7). Since low antibody responses are seen in humans infected or vaccinated with IBV, and ferrets inoculated with IBVs generate much lower immune responses compared to infection with seasonal IAVs, further investigation is required to identify ways to improve the immune responses to IBV.

Following exposure of the upper respiratory tract (URT) to influenza virus, the innate immune system is activated by interaction with pattern recognition receptors (PRRs) on susceptible cells. There are three different PRRs that sense influenza virus, which includes retinoic acid inducible gene I (RIG-I), Toll-like receptors (TLR3, TLR7 and TLR8) and the nucleotide binding oligomerization domain (NOD)-like receptors (8). Pro-inflammatory cytokines and type-I interferons are induced through RIG-I and TLR7 pathways (9). At the site of infection (respiratory epithelial cells), TLR3 and TLR7 sense viral double-stranded and single-stranded RNA (ssRNA) respectively within the endosome while RIG-I recognizes cytosolic ssRNA or viral RNA containing 5’-triphosphate and induces type I and III interferon (IFN) responses through the transcriptional factors NF-kB and IRF by interacting with MAVS. These interactions result in the activation of NF-kB and IRF3 which result in initiating a pro-inflammatory response by the production of cytokines and chemokines as well as an interferon type-I/III IFN response in respiratory epithelial cells. In IFN-knock out studies in mice, IFNB has been shown to play an important role in defense against IAV in lung epithelial cells, and IFNB absence results in delayed expression of IFNA thus, increasing the probability that the infecting virus can overrun the innate immune response of the host (10). The pro-inflammatory cytokines and chemokines can activate resident immune cells (innate lymphoid cells, macrophages, and dendritic cells) and recruit cells to the site of infection. Additionally, cytokines and chemokines can stimulate a T-helper 1 (TH1) response by IFNG and IL-12. They can also stimulate a T-effector response activation of STAT3 resulting in IL-6 and IL-23 secretion to further upregulate the inflammatory response (11). These cytokines bind to the epithelial cell IFN receptors (IFNAR) or those expressed by myeloid cells causing expression of interferon stimulating genes (ISGs) and an antiviral state. In parallel, cytosolic ssRNA and DAMPs (Danger-Associated Molecular Pattern) can also interact with NLRP3 of the inflammasome complex to cause cleavage and activation of caspase-1 and induction of IL-1b and IL-18. The regulatory effects on adaptive immunity are not a result of direct action of type-III IFN (IFNL) but rather indirectly through thymic stromal lymphopoietin (TSLP). IFNL secreted by infected respiratory epithelial cells can induce M cells to release TSLP which acts on migrating CD103+ dendritic cells (DCs) to stimulate the adaptive immune responses by enhancing CD8+ T-cell maturation and promoting germinal center reactions in the lymph node through T-follicular/helper (Tfh) cells (12). Tfh cells help in B-cell survival and proliferation in germinal centers (12–14) resulting in an increase of IgG1 and IgA antibody (14). Once these receptors are triggered by the virus, activation of several antiviral signals and production of cytokines and chemokines occurs to initiate the immune response in the host.

The humoral immune response plays a major role in immunity to influenza. This study focuses on the antibody response to influenza hemagglutinin (HA) which is the most commonly measured correlate of protection (15, 16). Cytokine and chemokine levels in serum have been shown to correlate with increased antibody responses following vaccination and infection. Interleukin-2 (IL-2) can work together with cyclic adenosine 3’, 5’-phosphate in synergy to enhance the antibody production by B-cells (17). In elderly human subjects, IL-2 treatment enhanced antibody responses to influenza vaccination (18). IL-2 has been shown to play an essential role in a T-helper 2 (TH2) differentiation (19) and may play a role in the adaptive immune response (20). Inflammatory cytokine levels of IL-6 and TNFA in serum have shown a positive correlation with fold-increases in hemagglutination inhibition titers following vaccination (21). IFN levels have also been shown to be protective and enhance antibody response following influenza infection. In mice lacking Tfh cells, which are important in humoral immunity, TH1 cells are the source of IFNG that promotes a protective IgG2 antibody response (22). IFNL has also been shown to increase adaptive mucosal antibodies (14). Taken together, this study explores innate and cytokine responses following IAV and IBV infection of ferrets and their contribution to development of robust adaptive antibody responses. Understanding factors which affect the ability of influenza viruses to generate robust immune responses may aid in the testing and improvement of influenza B vaccine efficacy in humans.

## RESULTS

### IAV virus replication kinetics differ from IBV and cause greater morbidity in ferrets

Ferrets were inoculated with influenza A/H1N1pdm09 (A/California/07/2009, “CA”), A/H3N2 (A/Kansas/14/2017. “KS”), B-Victoria lineage (B/Brisbane/60/2008, “BR”), or B-Yamagata lineage (B/Phuket/3073/2013, “PH”) virus Throughout the challenge, nasal wash and blood samples were collected for virus titration/molecular analyses and molecular/immunological analyses respectively.

Virus replication kinetics demonstrated peak early replication in the upper respiratory tract (URT) at D1 post challenge in IAV-challenged animals and late replication at D3 in IBV-challenged animals (Figure 1). IAV viral replication peaked earlier on D1 post challenge (CA=10^5.91^ FFU/mL, KS=10^5.30^ FFU/mL) whereas IBV virus replication peaked later at D3 post challenge (BR=10^5.24^ FFU/mL, PH=10^5.68^ FFU/mL) in ferret URT. There was a significant difference between IAV and IBV on D1 [CA vs BR (p<0.0001) and vs PH (p=0.007); KS vs BR (p=0.002)]; however, by D3 IBV replication increase to equivalent levels. PH, however, replicated to higher levels than KS on D3 (p=0.003). CA replicated significantly higher and earlier than IBV (BR, p<0.0001; PH, p=0.0236) as well as KS (p=0.0015). These replication kinetics confirm greater morbidity of CA in ferrets compared to other viruses tested and compliment previous findings of this virus compared to other seasonal influenza viruses (23).

**Figure 1:**
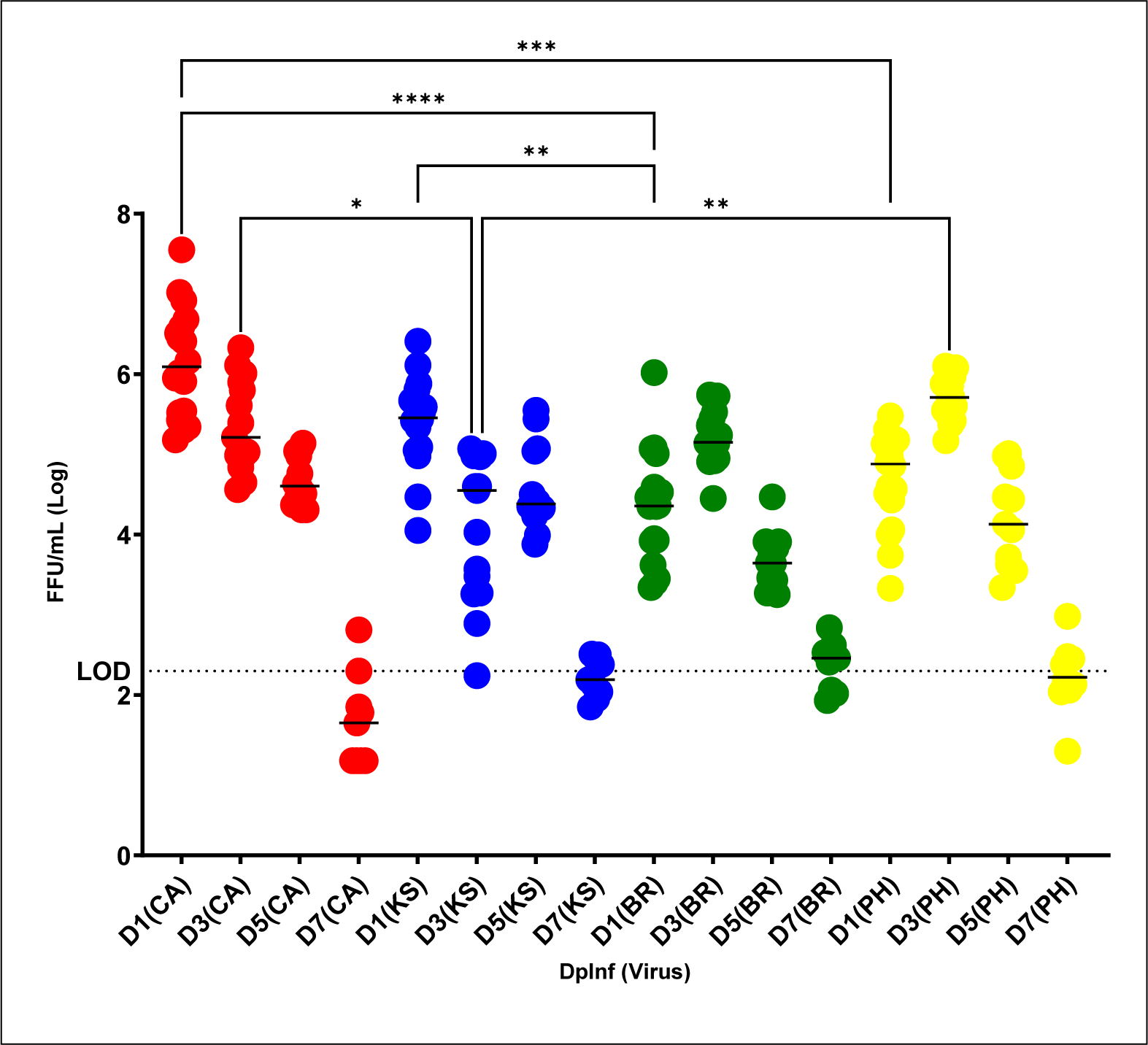
Viral kinetics of influenza in ferret upper respiratory tract. Replication kinetics of IAV [H1N1pdm09 “CA” (A/California/07/2009), H3N2 “KS” (A/Kansas/14/2017)] and IBV [B-Victoria lineage “BR” (B/Brisbane/60/2008), B-Yamagata lineage “PH” (B/Phuket/3073/2013)] in ferret upper respiratory tract. r 72Hr. A) H1N1pdm09 = red, H3N2= orange, B-Victoria = green, B-Yamagata= yellow. B) Comparison of IAV (CA/KS) and IBV (BR/PH) in black and white circles, respectively. All points are from four independent assays in FNEC conducted in triplicate. Limit of detection, indicated by a dotted line, was 10^3.3^ FFU/mL. Significance between groups was determined using 1-way ANOVA (Sidak’s multiple comparison test).

Following challenge with influenza viruses, ferrets exhibited classical clinical signs of infection including elevated temperature, weight loss, lethargy and sneezing as well as virus replication in the URT (Table 1). Regardless of the virus used, all animals developed fever at least +2°F over pre-challenge temperature by D2 post challenge with average temperature increases being similar between IAVs (+2.5°F) and IBVs (+2.55°F). No significant differences between groups were seen (2-way ANOVA). All animals demonstrated weight loss through D7 post challenge. The maximum weight loss from pre-challenge weight occurred on D3 for IBV [BR (-2.7%), PH (-4.9%)] and IAV [KS (-4.3%)]; however, CA peak weight loss occurred later on D6 (-7.6%). Significant weight loss was seen with IAV [CA (p<0.0001), KS (p=0.049)] compared to IBV (BR) over the first ten days following challenge (2-way ANOVA). Over the same period, PH weight loss was significantly greater than BR as well (p<0.0001). Daily temperature changes and weight loss can be found in supplemental Figure S1. For activity following challenge, only IAVs showed marked lethargy compared to IBVs ranging from IAV CA having the highest relative inactivity index (RII=1.37) followed by KS (RII=1.26) then the IBVs showing minimal lethargy BR (RII=1.04) and PH (RII=1.01). Taken together, clinical signs indicate that these IAVs cause greater morbidity in ferrets than the IBVs tested.

**Table 1:**
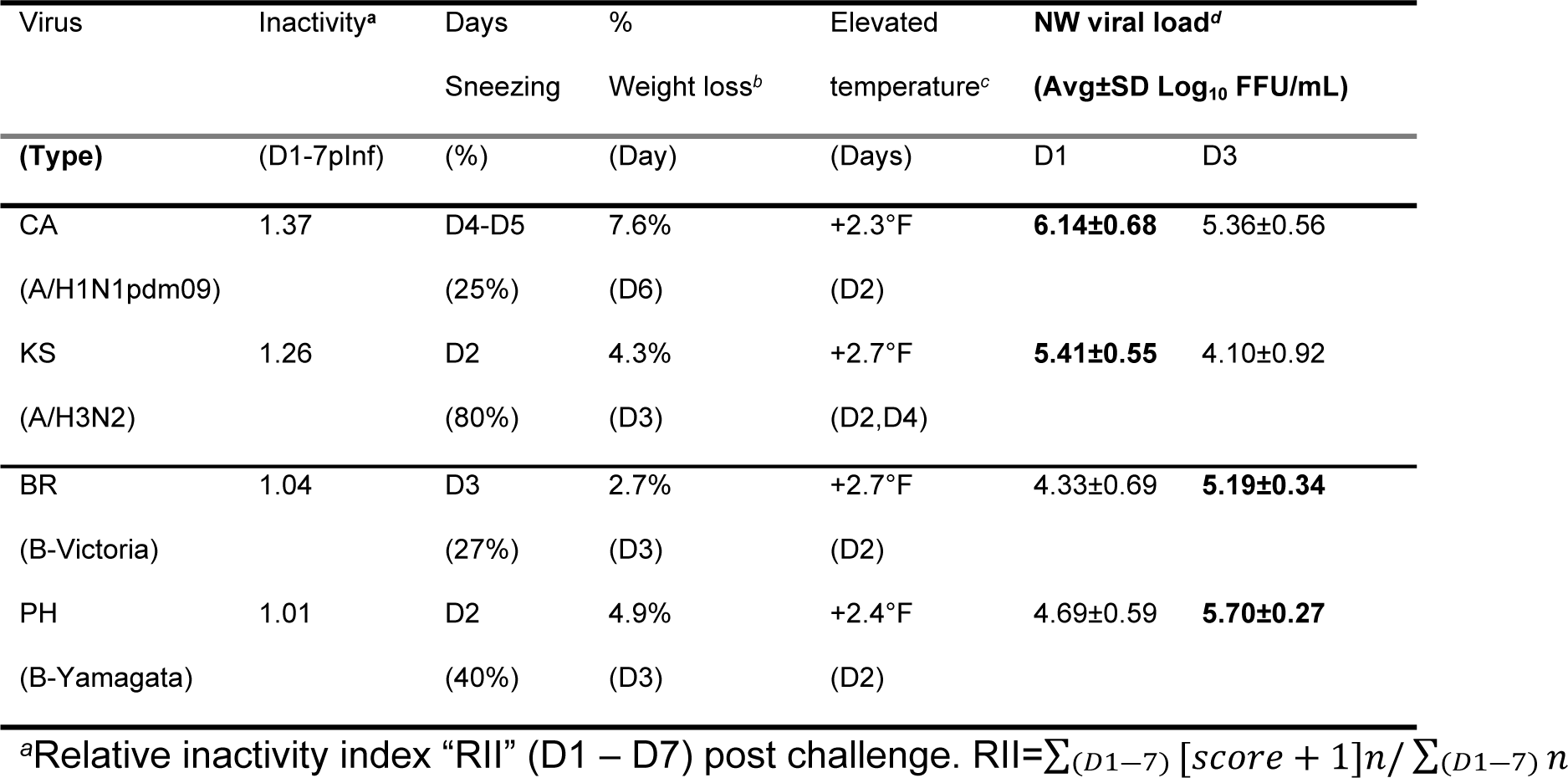
Ferret clinical signs following infection with influenza A and B viruses.

### IAV induces earlier inflammatory and interferon gene expression than IBV in ferret respiratory tract and peripheral blood

To determine the temporal relationship between innate immune responses and development of antibodies following challenge with influenza virus in ferrets, we conducted a comparison of host gene expression in NW cells in the URT and PBMC in the peripheral blood using qRT-PCR (Table 2). Gene expression at the site of infection in the URT (Table 2A) show that of the nine up-regulated genes (gene expression >1 over mock challenged) on D3 post challenge, seven genes (*MCP-1, CXCL10, STAT1, STAT2, STAT3, RIG-I, and SOCS3*) corresponded to either CA or KS infection. There were only two up-regulated genes(*TGFB1* and *TSLP*) following IBV (PH) infection. Of the remaining genes which were down-regulated on D3 (N=11) from pre-challenge expression, IBV down-regulated genes to a greater extent for all genes (7 of 11) except for *IL-1B, IL-6, IFNA, IFNL3*. By D5 post challenge four additional genes switched from down-regulated to up-regulated (*IL-2, IL-4, IFNG, IFNL3)* and were all following IAV (CA) challenge, indicating initiation of adaptive effector T cell and interferon responses by IAV.

**Table 2:**
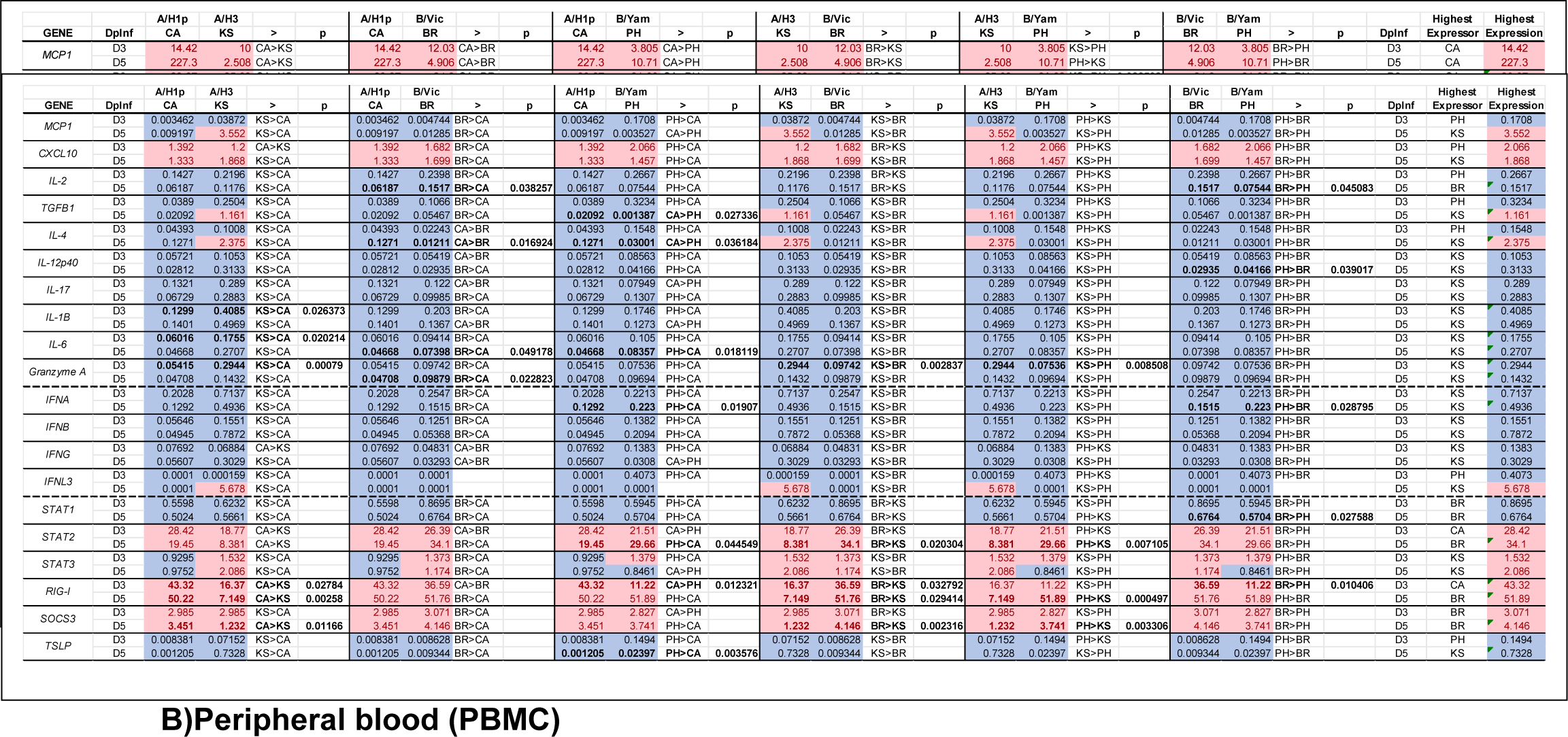
Comparison of ferret gene expression following IAV and IBV challenge in the URT and peripheral blood by qRT-PCR. **A)URT (NW)**

Gene expression in the peripheral blood (Table 2B) showed few up-regulated genes, compared to the URT, on both D3 and D5 post challenge. On D3 post challenge, five genes were up-regulated (*CXCL10, STAT2, STAT3, RIG-I, and SOCS3*). Three of these genes *(STAT2, STAT3, and RIG-I)* showed greater up-regulation following IAV infection; whereas two genes (*CXCL10 and SOCS3)* were up-regulated to a greater extent following IBV infection. Of the remaining genes which were down-regulated on D3 (N=14), IBV down-regulated genes to a greater extent for all genes (8 of 14) except for *IL-2, TGFB1, IL-4, IFNL3, STAT1 and TSLP*. By D5 post challenge, four additional genes were upregulated (*MCP-1, TGFB1, IL-4, and IFNL3)*. All of these genes were up regulated to the greatest extent by IAV. Of the original five genes which were up-regulated on D3, one gene switched from IBV to IAV (CXCL10) and two genes switched from IAV to IBV (*STAT2,* and *RIG-I*). These gene expression levels indicate that IAV resulted in an earlier innate immune response in both the URT and peripheral blood.

Comparison of gene expression by IAV and IBV in the (A) URT and (B) peripheral blood of ferrets on D3 and D5 post challenge by qRT-PCR. Gene expression in the URT was determined in NW cells and peripheral blood expression was determined in PBMC. Average fold-gene expression levels over pre-infection (D0) mock challenged animals for IAV (CA and KS) and IBV (BR and PH) are shown. An upregulation of expression (>1-fold) indicated in pink and a down-regulation (<1-fold) indicated in light blue. Comparisons of IAV to IAV (CA to KS), IAV to IBV (CA to BR, CA to PH, KS to BR, and KS to PH), and IBV to IBV (BR to PH) displayed and virus expression differences shown in column “>” and virus with highest level for all comparisons indicated in column “HIGHEST EXPRESSOR”. Comparisons with statistical differences, using t-test, p<0.05 shown in “p” column. Interferon genes are grouped between dotted lines.

### Serum cytokine and chemokine levels are elevated following IAV infection in ferrets

Serum cytokine and chemokine levels in ferrets following challenge with IAV and IBV were evaluated over a one-month period. Ferret sera were tested using a multiplex assay for ferret analytes (IFNB, IFNG, IP-10, IL-2, MCP-1, MIP-1B, IL-4, IL-17, IL-12p40, IL-12p70, TNFA, IL-6 and IL-8) and by bioassay (IFNL). Significant differences (2-way ANOVA) between IAV (CA and KS) and IBV (BR and PH) were observed for several analytes (Figure 2). Type -I (IFNB) and Type-II (IFNG) interferons showed significantly higher levels in ferret sera following challenge with IAV compared to IBV (Figure 2A). Type-I IFN levels were significantly higher for KS infected animals than CA (p=0.0152), BR, or PH (p<0.0001) infected animals. KS infected animals showed significantly higher levels of type-II IFN than CA (p=0.0401) as well as BR and PH (p=0.0405 and p=0.0431 respectively). Type-III IFN (IFNL) was higher in IAV than IBV-infected animals; however, only significant differences were found between BR and PH (p=0.0074) (Figure 2A). Additionally TH1 (IP-10), TH2 (IL-2) and T-eff (IL-12p40) cytokines (Figure 2B, 2C and 2D respectively) were also significantly higher following IAV than IBV challenge [IP-10; CA to KS, BR, and PH (p<0.0001) and KS to PH (P=0.004)], [IL-2; CA and KS to BR and PH (p<0.0001)] and [IL-12p40; CA and KS to BR and PH (p<0.0001)]. Pro-inflammatory cytokines and chemokines were also higher in IAV challenged than in IBV challenged animals. Pro-inflammatory response (Figure 2E) was significantly higher for: [MCP-1; CA to KS (p=0.0048) and BR and PH (p<0.0001)], [MIP-1B; KS to BR (p=0.0361)], and [TNFA; KS to BR (p=0.0468)]. Cytokine and chemokine levels over the entire 28-day period for each virus are shown in supplemental figure S2. Data from all analytes tested in sera show that for all groups (Pro-inflammatory, TH1, TH2, T-effector and IFN), IAV challenged resulted in higher levels than IBV challenged animals.

**Figure 2:**
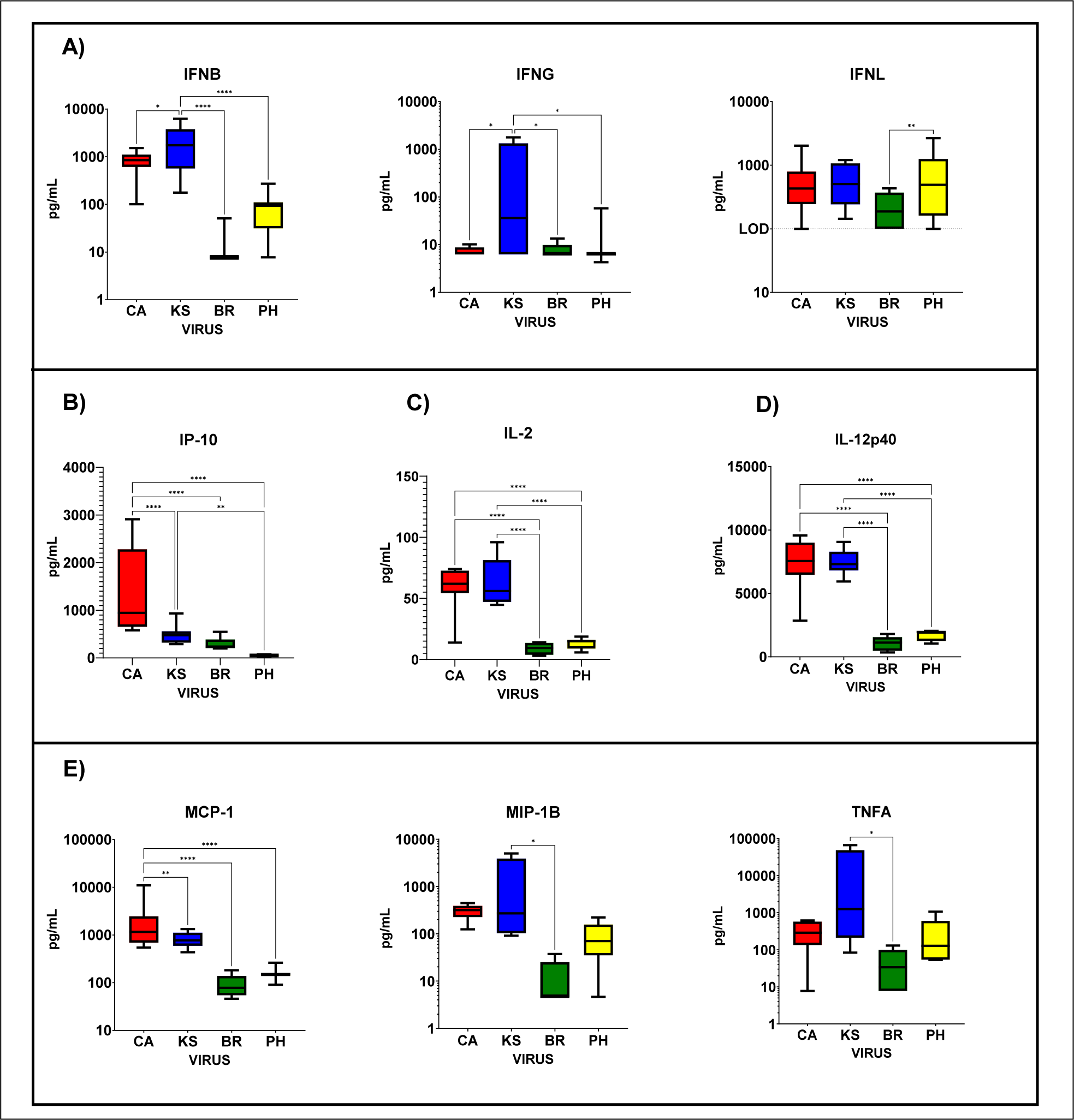
Cytokine and chemokine levels in ferret sera following IAV or IBV infection. Cytokine and chemokine levels in sera following IAV (CA and KS) or IBV (BR and PH) infection (day 1 to day 28) in ferrets. Influenza A viruses: H1N1pdm09 A/California/07/2009, “CA” (red) and H3N2 A/Kansas/14/2017, “KS” (blue). Influenza B viruses: B-Victoria lineage B/Brisbane/60/2008, “BR” (Green) and B-Yamagata lineage B/Phuket/3073/2013, “PH” (yellow). A) Type-I (IFNB), Type-II (IFNG) and Type-III (IFNL) interferons. B) TH1 chemokine (IP-10). C) TH2 cytokine (IL-2). D) T-effector cytokine (IL-12p40). E) Pro-inflammatory chemokines (MCP-1 and MIP-1B) and cytokine (TNFA). 2-way ANOVA used to determine significant differences between viruses over a 28-day period.

### Hemagglutination inhibition (HI) and neutralizing antibody responses are delayed and reduced in ferrets challenged with IBV compared to IAV

Serum antibody responses were compared between IAV and IBV challenged animals for 28-days. The antibody responses detected by HI (Figure 3A) showed clear differences between IAV (CA and KS) and IBV (BR and PH). For IAVs, infection with CA resulted in significantly greater antibody response than infection with KS (p=0.0094; 2-way ANOVA); however, for infection with IBVs no significant difference in antibody response was observed. Both IAVs showed significantly higher antibody responses to IBVs (CA vs BR/PH, p<0.0001; KS vs BR, p=0.0074 and KS vs PH, p=0.0005). By HI, sera from IAV surpassed the 1:160 threshold by D7 (CA, HI=2560) and D10 (KS, HI=640) post challenge; whereas only one IBV (BR) surpassed the threshold but not until D28 post challenge (HI=201). Neutralizing antibody responses by FRA (Figure 3B) mimicked antibody responses seen by HI; however, due to its greater sensitivity at lower antibody concentrations (Figure 3C), both IBVs crossed the 1:160 threshold but not until very late post challenge (BR on D21, FRA=320; PH on D28, FRA=186). Both IAV showed significantly higher neutralization titers compared to IBV for all viruses (p<0.0001); however no statistical differences between IBV (BR and PH) were observed. Results of antibody responses in ferrets showed that IAV initiated an earlier and higher neutralizing antibody titer that was sustained over the 28-day period. In contrast, IBV neutralizing antibody titers were delayed and failed to reach comparable titers to IAV by D28. Both HI and neutralizing antibody (FRA) responses in ferrets following IAV or IBV infection are consistent with antibody responses seen with previously tested viruses.

**Figure 3:**
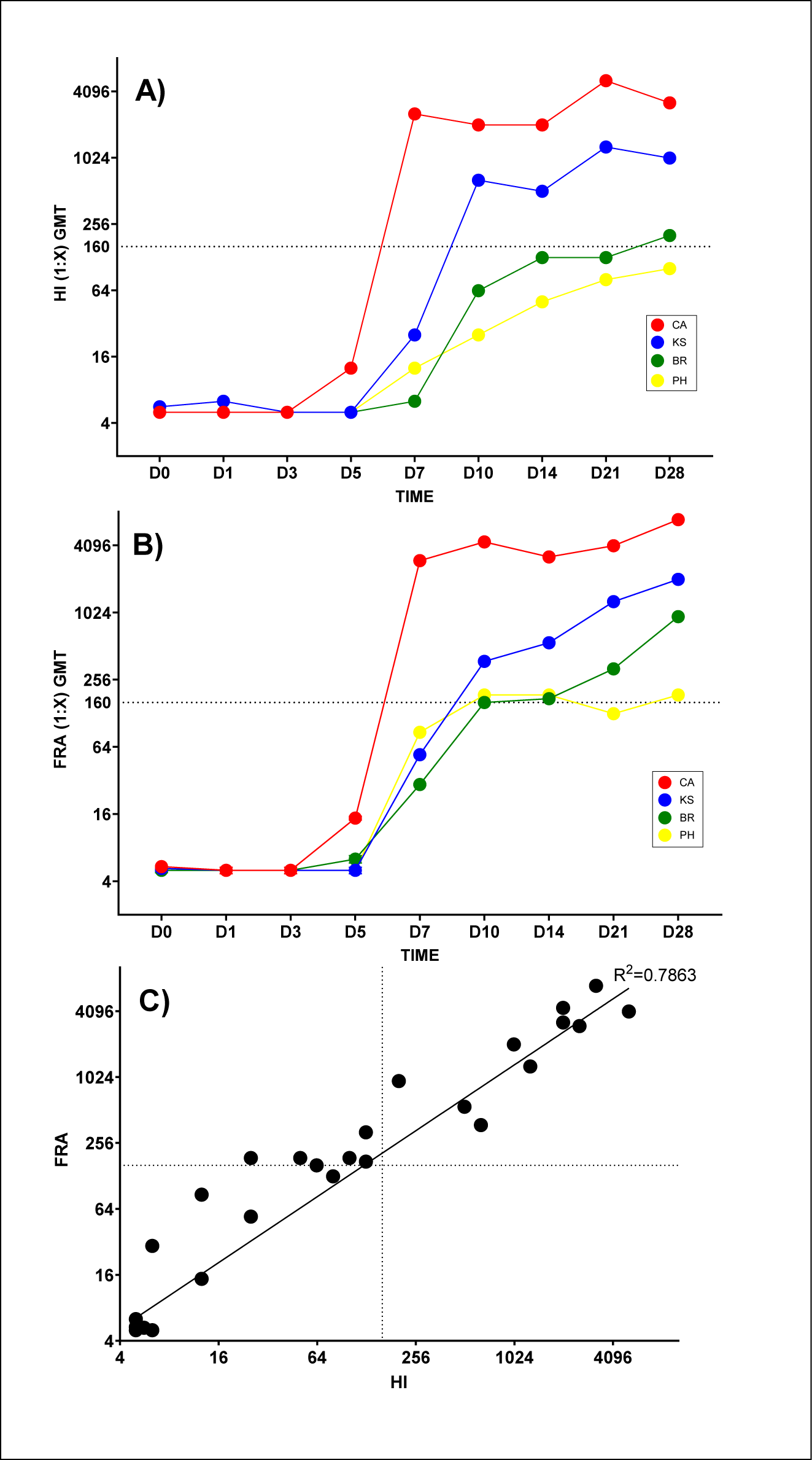
Serum antibody responses following IAV or IBV ferret challenge. Serum antibody by HI and neutralizing antibody (FRA) levels following infection with IAV (“CA” red, “KS” blue) and IBV (“BR” green, “PH” yellow) over 28 days post challenge. Average geometric mean titer (GMT) given for HI (A) and FRA (B) tests. Values above 1:160 (dotted line) indicate robust antibody response. Correlation of HI to FRA (C).

To evaluate the kinetics and magnitude of the ferret Ig response elicited following IAV or IBV infection, sera collected throughout the study were assessed for reactivity against the corresponding rHA protein by ELISA (Figure 4). Consistent with a delay in the generation of IgG class-switched antibodies in the primary response, increased rHA-specific antibody titers were not robustly detected until Day 10 for both IAV and IBV. For antigen-specific ELISA to IgG (Figure 4A), similar antibody patterns were observed for each individual virus. When comparing IgG responses for IAV and IBV (Figure 4C), IBV showed significantly higher levels on D14 (p=0.034); however, by D21 antibody IgG titers to IAVs were significantly higher than to IBVs (D21 p<0.0001, D28 p=0.0001). Interestingly, when performing antigen-specific ELISA for total Ig (IgKappa and IgLambda light chains) “totIg”, individual IBVs showed higher titers than individual IAV over 28-days post challenge (Figure 4B). When comparing IAVs to IBVs (Figure 4D), significantly higher levels of totIg were observed following IBV infection starting on D10 through D28 compared to IAV infection (D10/D14 p<0.0001; D21 p=0.0011; D28 p=0.0003).

**Figure 4:**
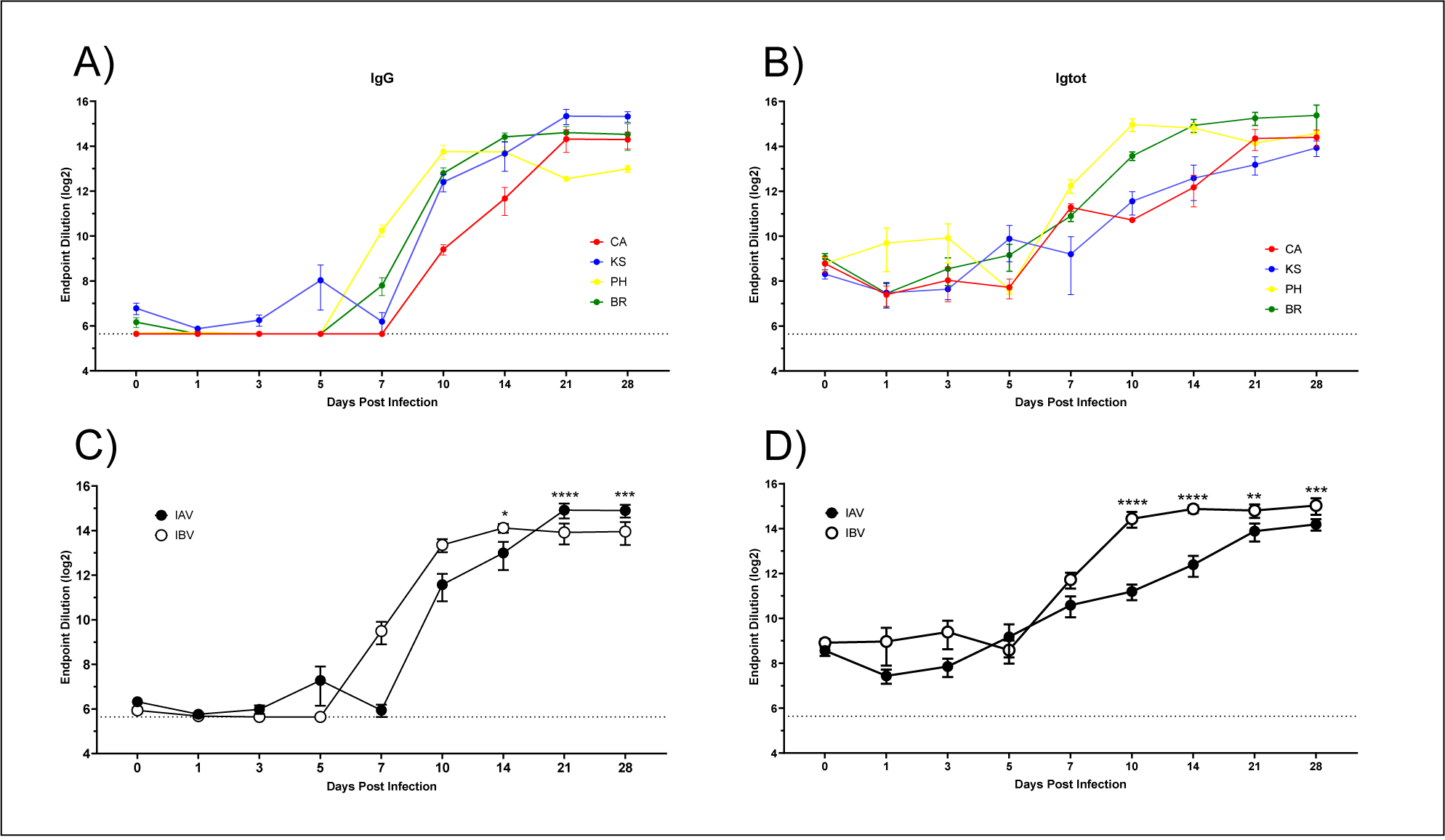
Antigen-specific antibodies in ferret serum following IAV and IBV challenge by ELISA. Antigen-specific antibodies in ferret sera over 28 days following infection with IAV and IBV. Antigen-specific IgG (A) and total Ig (B) to CA (red), KS (blue), BR (green) or PH (yellow) over 28 days. Comparison of IAV to IBV for antigen-specific IgG (C) and total Ig (D). Significant differences for each timepoint determined using 2-way ANOVA. Dotted line indicates limit of detection.

### Delay in serum antibody responses by IBV are not observed by antibody secreting cells

To determine whether the delay and reduced antibody levels in serum were also observed in antibody secreting B cells, we performed ELISpot assays to detect ferret IgG or total Ig “totIg” (IgKappa and IgLambda light chains) (24) antibody secreting cells (ASC) in ferret splenocytes or PBMC on D5 and D7 post IAV and IBV challenge (Figure 5). In ferret splenocytes (Figure 5A and 5B) significant increases on D7 in ASC secreting IgG and totIg in IBVs (BR and PH) and IAV (KS) were detected [(IgG: BR to KS, p=0.0038; PH to KS, p=0.047); (totIg: BR to KS, p=0.008; PH to KS, p=0.044)]; however no significant increases of IBVs to IAV (CA) were detected. When comparing differences in both IAV to both IBV, no significant differences were detected. Similar responses were observed in ASC from PBMC (Figure 5C and 5D) comparing IBV (PH) to IAV (KS). Even though some differences in ASC were observed with individual viruses, overall differences between IAV and IBV were absent. This is in contrast to significant differences seen between serum antibody responses by HI and FRA where IAV virus antibody responses appeared earlier and more robust than IBV following challenge. While IAV elicits potently neutralizing antibody specificities, IBV may elicit strong rHA-specific response without potent HI or FRA activity. This may be due to differences in epitope bias.

**Figure 5:**
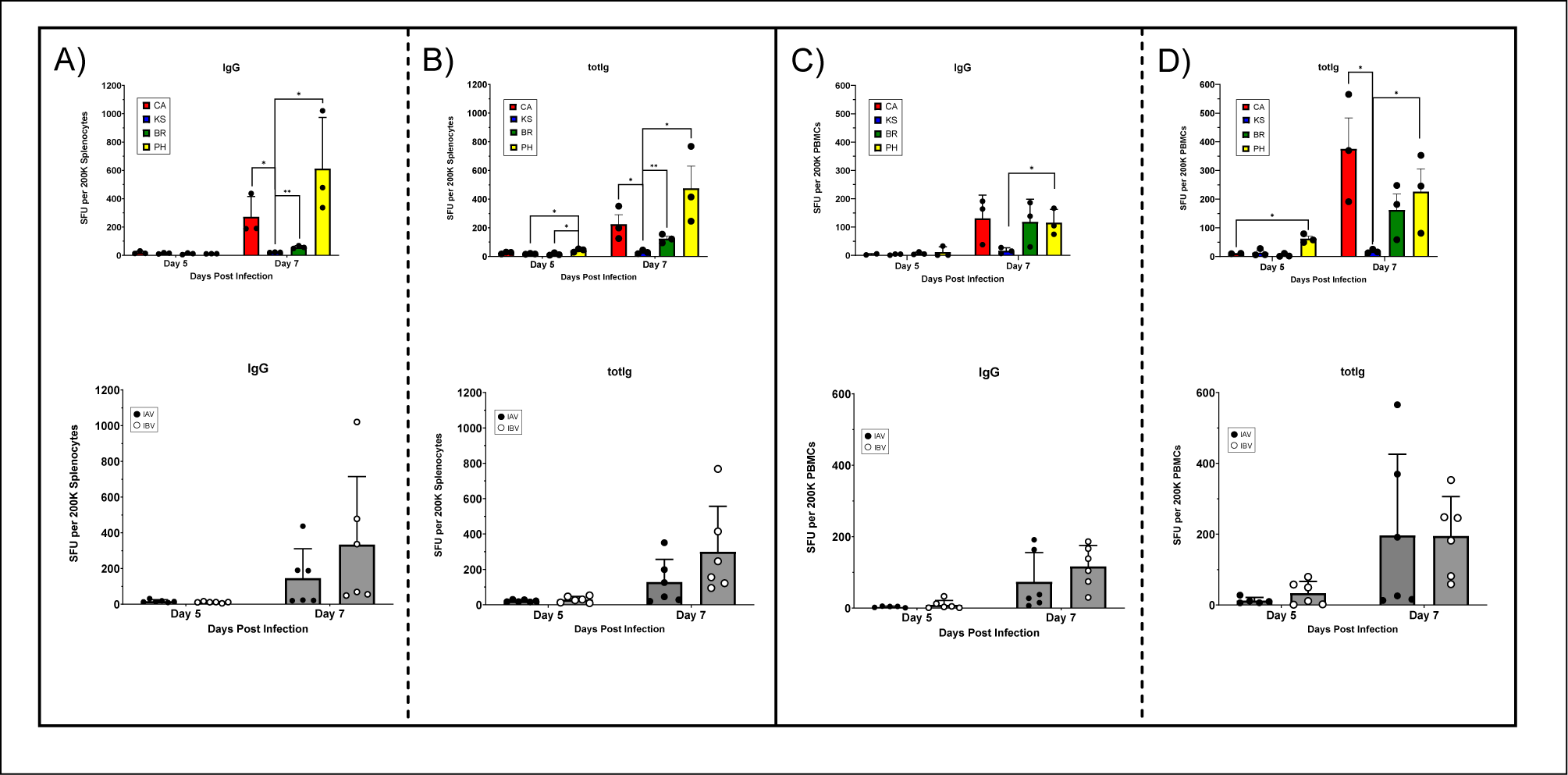
Antibody-secreting cells in ferret spleen and peripheral blood following IAV and IBV challenge. Early antibody-secreting cells (ASC) in splenocytes (A and B) and peripheral blood (PBMC) (C and D) on days 5 and 7 post challenge with IAV and IBV. ASC to following challenge with individual viruses: CA (red), KS (blue), BR (green) or PH (yellow) in splenocytes and PBMC. Panel A) top panel is individual virus IgG spot forming units (SFU) in splenocytes (significance using multiple t-tests), bottom panel comparing IgG SFU following IAV and IBV challenge (ns, t-test). Panel B) top panel is individual virus total Ig “totIg” SFU in splenocytes (significance by multiple t-tests), bottom panel comparing totIg SFU following IAV and IBV challenge (ns, t-test). Panel C) top panel is individual virus IgG SFU in PBMC (significance by multiple t-tests), bottom panel comparing IgG SFU following IAV and IBV challenge (ns, t-test). Panel D) top panel is individual virus total totIg SFU in PBMC (significance by multiple t-tests), bottom panel comparing totIg SFU following IAV and IBV challenge (ns, t-test).

### Correlations between viral loads, gene expression, antibody responses and protein levels in ferret samples

We compared virus levels in the URT (D1, D3, and D5 post challenge) to gene expression in both cells in the NW of the URT and PBMC in the peripheral blood (Table 3) to determine whether there was a correlation of viral titers to innate gene expression in IAVs and IBVs. In the URT (Table 3A) we saw a direct correlation between type-I IFN (*IFNA, IFNB*) and gene expression and IAV viral titers; however, there was an inverse correlation with IBV. Similar correlations were also seen with T-effector gene (*IL-17*) and viral titers. For TH1 (*CXCL10*) we saw the reverse; a direct correlation for IBVs and an inverse correlation for IAVs. Generally, we saw greater direct correlations between viral titers and IFN and pro-inflammatory genes with IAV (KS) than other viruses. Interestingly a direct correlation between both *IFNL* and *TSLP* gene expression and viral titers was observed for both IAV (KS) and IBV (PH) indicating that IFNL and TSLP gene expression may be linked. However, only inverse correlations for inflammatory gene (*MCP-1)* and interferon response gene (*STAT2)* showed any significance in IBV (PH). For correlations between viral titers in the URT and gene expression in PBMC (Table 3B), a direct correlation was achieved for type-I/II IFN (*IFNA, IFNB, IFNG)* genes in all IAVs and IBVs (*IFNA*) and three of four viruses (*IFNB* – CA, KS, and BR; *IFNG* – CA, KS, and PH). A direct correlation of pro-inflammatory gene (*IL-1B*) to virus titer was also seen in all IAVs and IBVs. An inverse correlation was achieved for IFN-response genes (*STAT2* and *RIG-I*) and all IAVs and IBVs indicating that direct effects on IFN expression may result in lower virus titers. Even though only a few comparisons showed significant correlations, due to a limited number of observations (D1, D3, D5), overall comparisons of URT virus titers and gene expression in NW cells and PBMC showed a trend that IAVs impacted more genes with a direct correlation to pro-inflammatory and IFN responses than IBVs.

**Table 3:**
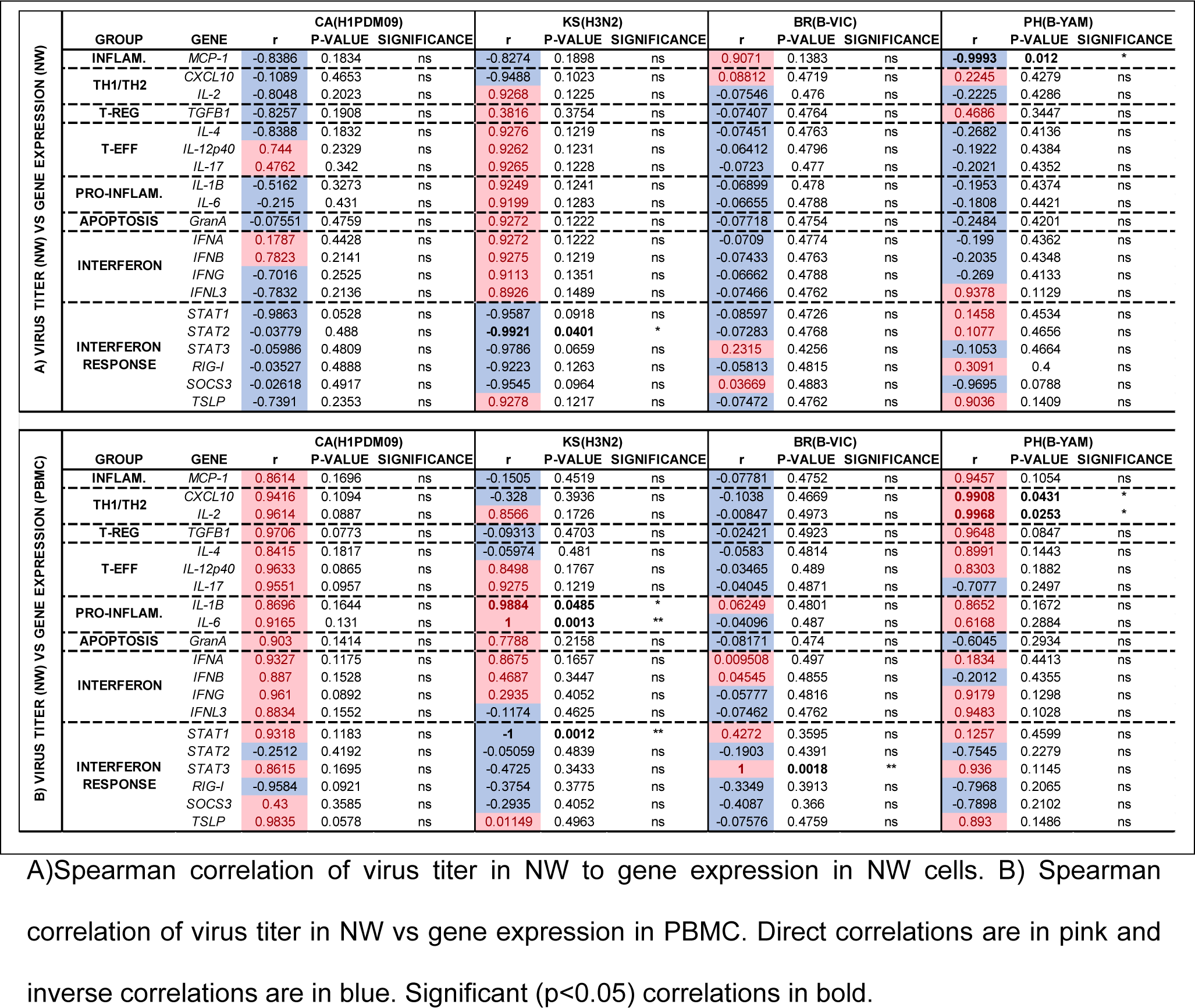
Correlation between virus titer and gene expression in ferret samples.

Since additional timepoints were tested (pre-challenge and days 1, 3, 5, 7, 10, 14, 21 and 28 post challenge), a clearer correlation was seen when comparing antibody levels (HI and FRA) to analyte levels in ferret sera following IAV and IBV challenge (Table 4). Direct correlations of serum HI (Table 4A) and neutralizing FRA (Table 4B) antibody was seen for all IAVs and IBVs to TH2 (IL-2) and T-effector (IL12-p40) analytes. These analytes indicated that strong adaptive (TH2 and T-effector) responses were associated with increased antibody responses. Interestingly, an inverse correlation between the pro-inflammatory response (MCP-1 chemokine) was observed for all IAVs and IBVs indicating that a strong pro-inflammatory response may dampen the antibody response. Some differences in correlation are seen comparing HI and FRA to analyte levels. For instance, IFNB showed a direct correlation to HI antibody response to all IAVs and IBVs (Table 4A); however, by FRA, only IAV (KS) and IBV (PH) showed a direct correlation (Table 4B). Also, T-effector (IL-4) showed an inverse correlation to all IAVs and IBVs by FRA (Table 4B); however, this correlation was only observed for 3 of 4 viruses by HI (Table 4A). Correlations to IFNs are also seen depending on whether HI or FRA is used for the comparisons. For example, type-III IFN (IFNL) levels showed direct correlation to IAV and inverse correlation to IBV using HI (Table 4BA) ; however, by using FRA, this is not a clear distinction (Table 4B). Since analytes are secreted with different kinetics over the entire 28 days (Supplemental Figure S2), a clearer distinction could be obtained if the window of comparisons were narrowed. Overall differences in the analytes expression, as well as gene expression, showed differences in immune responses to IAVs and IBVs.

**Table 4:**
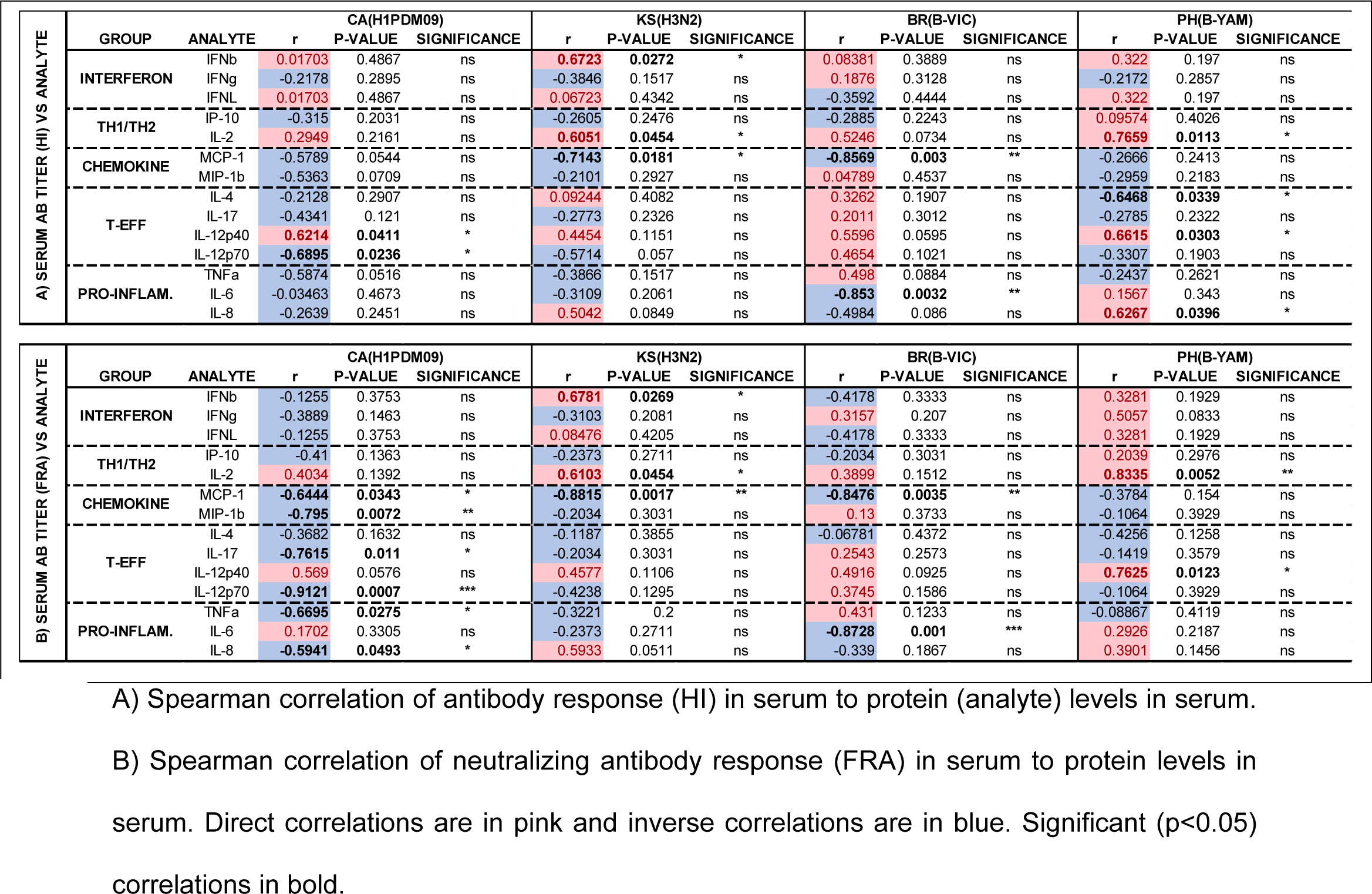
Correlations between antibody levels and protein levels in ferret sera.

## DISCUSSION

When using antisera generated in ferrets following challenge with influenza viruses, we noted that while IAVs generated robust antibody responses fourteen days post challenge, IBV challenge resulted in delayed and significantly lower antibody responses (HI titers <1:160) over the same period in almost 90% of the viruses tested. This study was designed to determine whether innate immune markers and soluble cytokines/chemokines in serum following challenge could shed light on why IBVs do not elicit an antibody response as quickly and robustly as IAVs in ferrets.

Four representative viruses were used in this study, two IAVs and two IBVs to determine innate and adaptive differences following challenge in ferrets. Initial differences in clinical signs as well as virus kinetics following IAV or IBV are apparent. Clinically, IAVs caused greater lethargy (average RII=1.31) compared to IBV (average RII=1.03), greater maximum weight loss (IAV ∼6% and IBV ∼4%), and more sneezing (IAV ∼52%; IBV ∼33%) during viremia. In addition to greater morbidity, IAVs virus replication peaked early (D1 post challenge) compared to IBV (D3 post challenge) in ferrets. These observations could indicate that IAVs initiate an earlier innate immune response than IBVs which could lead to an earlier and more robust adaptive immune response. It has been shown for COVID-19 vaccination that a strong innate immune response results in an early, robust antibody response(25).

We have previously shown in ferret primary respiratory tract cells that pro-inflammatory genes and IFNs are delayed and downregulated by IBVs compared to IAVs (26). TH1 (*CXCL10*), TH2 (*IL-2*), and T-effector (*IL-4*) were also significantly up-regulated by IAV. Similar responses were seen in the URT and PBMC from IAV and IBV infected ferrets in this study. IAV (CA), in particular showed early (D3) up-regulation of TH1 (CXCL10), and late (D5) up-regulation of TH2 (*IL-2*) and T-effector (*IL-4*) in the URT indicating a switch to a strong adaptive immune response. For IFN response in the URT, only Type-II (*IFNG*) and Type-III (*IFNL3*) IFNs were up-regulated from pre-challenge levels following IAV (CA) challenge. In PBMC by D5 post challenge only T-effector (*IL-4*) and Type-III IFN *(IFNL3*) were up-regulated by IAV (KS). However, early D3 up-regulation TH1 (*CXCL10*) was highest for IBV (PH) which switched to late D5 highest up-regulation for IAV (KS). Interestingly IFN response (*SOCS3*) gene, which is responsible for downregulating the IFN response was highest for IBV (BR) both early and late post challenge. This may support that IBV have greater IFN-antagonism properties than IAV. Unfortunately, due to the limited number of samples tested (3 animals per timepoint) significant differences for gene expression for all genes was not possible. Additional studies with greater samples collected and tested at each timepoint may aid in increasing the significance as was seen in our study using ferret respiratory epithelial cells. However, it is important to note that the responses seen in this study showed similar trends as observations from respiratory epithelial cells infected with IAV and IBV.

Striking differences in antibody responses to IAVs and IBVs were observed that indicated differences in the initial innate immune response may be playing an essential role in triggering a strong adaptive immune response. In both HI and neutralizing antibody tests, IAV infections generated an early, strong antibody response between day 7 to 10 post challenge, whereas IBV infections did not reach equivalent HI titers and required at least 21 to 28 days to reach titers of at least 1:160 by FRA. These experiments terminated at 28 days post challenge; it would be interesting to see if IBV antibody responses eventually reach equivalent levels to IAVs after 1 month or if peak antibody levels are reached by 28 days post challenge. Additionally, it would be important to determine how long IAV antibody titers remain elevated above the 1:160 threshold. Antibodies to influenza A virus may persist for up 18 months or longer in humans (27–29), however, the longevity of influenza B virus directed antibodies has not been tested.

Even though significant suppression and delay in serum HI and neutralizing antibodies was observed, antibody responses to recombinant antigens by ELISA showed significantly higher total Ig antibodies and earlier IgG antibodies to IBV compared to IAV. Similar differences in IgG responses by ELISA compared to HI and neutralizing titers were seen by Huang *et al.* (30). As observed in this study, a comparison of thirteen IAV/IBV strains in ferrets by Huang *et al.* showed that IBV induced the weakest antibody response. Thus, a direct determination of Ig antibody levels by ELISA may not be sufficient in the development of a strong antibody response. Additionally, the four canonical antigenic sites may be detected differently by HI/FRA and ELISA. It has been demonstrated in a study using IBV (B/Yamagata/16/1988) that the HA 120 loop appears to be the least dominant to elicit HI antibodies in mice and ferrets but contributed significantly to the total IgG responses by ELISA; conversely, the 190 helix induced substantial HI antibodies but the least dominant to elicit total IgG responses in the ferret (31).

Comparisons of ASCs in ferret splenocytes and PBMC following IAV and IBV infection revealed that although there were significant differences between antibody responses in the serum, these differences were not seen at the B-cell level. We observed the highest number of IgG and total Ig ASCs following IBV (PH) challenge and the least number of ASC following IAV (KS) challenge on day 7. IAV (CA) however, showed equivalent levels of IgG and total Ig ASC in PBMC. When comparing all IAVs (CA and KS) to all IBVs (BR and PH) there were no statistical differences in either splenocytes or PBMC. Greater differences between IAV and IBV may be seen in ASC in the mediastinal lymph nodes (32).

To determine whether there were differences in cytokine and chemokines levels in ferrets following IAV and IBV infection over 28 days, we tested sera by multi-plex or bioassay (IFNL). IAV infection resulted in significantly higher levels of Type-I/II IFNs, TH1 (IP-10), TH2 (IL-2), T-effector (IL-12p40) and pro-inflammatory cytokines/chemokines (MCP-1, MIP-1B, and TNFA). These responses showed correlations to increases in antibody responses detected by HI and FRA. These responses indicate that strong adaptive responses are associated with robust antibody responses. This further supports our assumption that a strong innate immune response will lead to a robust antibody response.

In conclusion, the data suggest that IBVs have a greater suppression of the innate immune response than IAV and this would lead to a reduced antibody response. In this report, inflammatory and IFN gene expression is higher following IAV than IBV infection in the URT and PBMC of ferrets. Additionally, serum levels of pro-inflammatory cytokines and type I/II IFNs are significantly higher following IAV infection than IBV infection. Suppression of the IFN response is a hallmark of immune evasion by influenza (33). Further studies adding immunoadjuvants, such as type-I or type-III IFNs to vaccines or following infection may aid in overcoming innate immunosuppression by IBVs resulting in generation of robust antibody titers leading to better protection. This may be important and beneficial in developing improved vaccines for influenza and other viruses of public health importance.

## MATERIALS AND METHODS

### Viruses

Influenza viruses used were passaged in Madin Darby Canine Kidney (MDCK) cells to retain antigenic similarity to the original human isolates and to avoid structural changes in the hemagglutinin which could occur from passage in eggs (34–36). All influenza viruses were passaged in MDCK “C#” or MDCK-SIAT1 ”S#” (37) cells according to established procedures (38, 39). Representative influenza A viruses used in this study were: A/California/07/2009 “CA” (H1PDM09 subtype; passage C3) and A/Kansas/14/2017 “KS” (H3N2 subtype; passage S3). Representative influenza B viruses used in this study were: B/Brisbane/60/2008 “BR” (B-Victoria lineage; passage CXC6) and B/Phuket/3073/2013 “PH” (B-Yamagata lineage; passage C4). Viral titers were determined by focus forming assay (26) and given as log Focus Forming Units per milliliter (FFU/mL). Viral titers for virus stocks used in this study were: CA (10^6.64^ FFU/mL), KS (10^7.00^ FFU/mL), BR (10^7.17^ FFU/mL) and PH (10^7.54^ FFU/mL).

### Ferret infection with influenza viruses and monitoring

All animal procedures were approved by the Institutional Animal Care and Use Committee of the Centers for Disease Control and Prevention in an Association for Assessment and Accreditation of Laboratory Animal Care International accredited facility. Male fitch ferrets (Triple F Farms, Sayre, PA), between 10 to 18 months of age and seronegative for currently circulating influenza viruses were used for infection experiments.

Four separate experiments were conducted using 18 ferrets for each virus [CA (influenza A/H1PDM09), KS (influenza A/H3N2), BR (influenza B/Victoria lineage) and PH (influenza B/Yamagata lineage)]. In each experiment ferrets were first anesthetized with an intramuscular cocktail of ketamine (15-30 mg/kg) plus xylazine (1-2 mg/kg) “KX” and challenged intranasally with 1 mL of virus (4×10^5^ FFU/mL). This dose of virus has been optimized and used as the standard dose for generating antiserum to human influenza clinical isolates in ferrets. The animals were monitored over 28 days post challenge for weight loss, fever, and other clinical signs (lethargy, sneezing and dyspnea). The animals were weighed on day 0 prior to influenza challenge to establish a baseline weight and then weighed daily for the first ten days post challenge and followed weekly thereafter through day 28. This corresponds to typical viral kinetics observed in the upper respiratory tract of ferrets infected with seasonal influenza viruses (40, 41). Changes in weight were calculated as percentage loss or gain from day 0 weight. For fever calculation, a temperature transponder [IPTT-300; Bio Medic Data Systems (BMDS), Waterford, WI, USA] was programed to identify each ferret and implanted subcutaneously between the shoulder blades of each animal and read with a scanner (DAS-6007 IPTT Scanner System, BMDS). A baseline temperature (°F) was determined by the average temperature over three days prior to challenge. Temperatures were assessed over the first 10 days post challenge, then weekly and fever was determined as any temperature greater or equal to 2 degrees Fahrenheit above baseline. Ferrets were assessed daily for clinical signs within a two-hour window between 9AM – 11AM during peak normal activity and to reduce variation due to circadian rhythms (42). Lethargy was determined by the relative inactivity index (RII) from day 0 through day 7 post challenge; this period allowed for lethargy calculation throughout the period of active replication in the ferret respiratory tract. RII, sneezing and dyspnea were assessed each day prior to handling and sedation of the animals. Clinical signs assessment was consistent with previously established methods (43, 44).

### Ferret sample collection and processing

Pre-challenge and on days 1, 3, 5, 7, 10, 14, 21 and 28 post challenge, nasal wash (NW) and blood samples were collected from ferrets for virologic, genetic and antibody analysis. The animals were monitored for clinical symptoms of influenza virus infection and lethargy.for seven days post infection. This corresponds to typical viral kinetics observed in the upper respiratory tract of ferrets infected with seasonal influenza viruses (40, 41). Animals were sedated with KX followed by flushing the nares by instillation of 2 mL nasal wash solution (PBS, 1% BSA, antibiotics), inside a class-II biosafety cabinet, and collected when expelled in a Petri dish. NW was centrifuged (5 minutes x 5000 rpm at 4°C), supernatant was stored at -80°C for virus titration and multiplex assay. One hundred microliters (uL) PBS was added to NW pellets followed by 280uL AVL lysis buffer (Qiagen, Germantown, MD, USA). Two to three mL blood was collected from the cranial vena cava in an SST tube for serum separation. Serum was tested for antibodies to influenza virus, cytokines, and chemokines.

On days 1, 3, 5, 7, 10 and 28 post challenge, three ferrets were euthanized for collection of tissues, sera, and cells. These animals were first anesthetized with KX, exsanguinated, and euthanized with intracardiac injection of Euthasol™ (1 mL/kg) prior to collection of tissues, including spleen. Blood was collected in SST tubes (serum) and K2/EDTA (cell purification). Serum was separated from SST tubes by centrifugation for 10 minutes at 1500 x g at room temperature. For purification of peripheral blood mononuclear cells (PBMC), an equal volume of PBS was added to blood in K2/EDTA tubes and overlayed onto Histopaque®-1077 (Sigma, St. Louis, MO, USA). Samples were centrifuged with no brake at 900 x g for 20 minutes at room temperature. Cells were collected from the Histopaque® interface, washed with cell culture media (RPMI-1640, 10% heat-inactivated fetal bovine serum (FBS)) and treated with ammonium chloride to lyse any remaining red blood cells. The final cell pellet was resuspended in 1 mL cell culture medium. Two hundred eighty uL of AVL lysis buffer were added to 200 uL PBMC samples in RPMI-1640, 10% FBS obtained by density gradient centrifugation over ficoll-hypaque. Samples were mixed 3-4 times and lysate was frozen at -80°C until RNA extraction. Carrier RNA was added to each sample and RNA was extracted using EZ1 DSP kit on a Qiagen EZ1 Advanced XL extractor according to the manufacturer’s instructions (Qiagen, Germantown, MD, USA). RNA was eluted in 120 uL RNase-free water and stored at -80°C until evaluated for gene expression by qRT-PCR. The remaining PBMC sample was cryopreserved in FBS, 10% DMSO and stored in the vapor phase of liquid nitrogen freezer. Splenocytes were isolated from spleens by passing tissue through a 70-micron cell strainer (Millipore-Sigma, Burlington, MA, USA) in DMEM,10% FBS, followed by lysing red blood cells with ammonium chloride. Cells were cryopreserved in 10% dimethyl sulfoxide, 90% cell culture medium and stored in the vapor phase of liquid nitrogen freezer until used in ELISpot assays.

Tissues and blood were collected from an additional three mock-infected ferrets and included as naïve controls.

### Virus Kinetics by focus forming assay (FFA)

Nasal wash (NW) samples collected from influenza virus-infected ferrets were tested for virus with a Focus Forming Assay (FFA) as previously described (26). Briefly, ½-log serially diluted influenza viruses in virus growth medium plus trypsin (DMEM, 0.1% BSA, 1 µg/mL TPCK-treated trypsin (Sigma, St. Louis, MO, USA)) were added to confluent monolayers of MDCK-SIAT1 cells (37) in 96-well flat-bottom tissue culture plates in quadruplicate. Following a two-hour incubation at 37°C, and overlay of 1.2% Avicel RC/CL (45) (Type: RC581 NF; FMC Health and Nutrition, Philadelphia, PA, USA) in 2X MEM containing 1 µg/mL TPCK-treated trypsin, 0.1% BSA, and antibiotics] was added. Plates were incubated overnight at 37**°**C, 5% CO_2_, fixed, permeabilized, and stained with a monoclonal antibody pool to influenza A or B nucleoprotein (International Reagent Resource; www.internationalreagentresource.org). Infectious foci (spots) were visualized using TrueBlue substrate (Sera Care, Inc., Milford, MA, USA). The foci were enumerated using a CTL Bio Spot Analyser with ImmunoCapture 6.4.87 software (Cellular Technology Ltd., Shaker Heights, OH, USA). The FFA titer was determined by multiplying sample dilution which gave between one hundred to three hundred spots by the spot number at that dilution, to obtain the Focus Forming Units per milliliter (FFU/mL). The foci in the cell control were subtracted and the number of foci remaining was multiplied by twenty to give FFU/mL. The limit of detection was 10^2.3^ FFU/mL.

### Antibody detection by hemagglutination inhibition (HI) assay

Ferret sera were treated with receptor destroying enzyme [RDE] (Denka Co Ltd, Tokyo, Japan) and adsorbed with turkey erythrocytes (TRBC), and tested by HI assay according to established procedures (38). Briefly, RDE-treated and adsorbed sera were 2-fold serially diluted, starting at 1:10, in v-bottom 96-well plates followed by the addition of 4 hemagglutination units of influenza virus (CA, KS, BR, or PH). Plates were incubated at room temperature for approximately 15 minutes followed by the addition of 0.5% TRBC and mixed by gentle agitation. The TRBCs were allowed to settle for 30 minutes at room temperature. The HI titers were determined as the reciprocal of the last serum dilution which inhibited the hemagglutination of the TRBCs by the virus.

### Neutralizing antibody level detection by focus reduction assay (FRA)

The Focus Reduction Assay (FRA), initially developed by the WHO collaborating Centre in London, United Kingdom, was modified and utilized in this study. MDCK-SIAT1 cells were seeded in 96-well plates in Dulbecco’s Modified Eagle Medium (DMEM) containing 5% heat-inactivated FBS and antibiotics. The following day, the confluent cell monolayers were rinsed with 0.01M phosphate-buffered saline pH 7.2 (PBS), (Gibco BRL, ThermoFisher Scientific Inc., Waltham, MA, USA) followed by the addition of two-fold serially diluted RDE-treated ferret sera at 50 µL per well starting with 1:20 dilution in virus growth medium containing 1 µg/mL TPCK-treated trypsin, VGM-T [DMEM, 0.1% fraction-V bovine serum albumin (BSA), antibiotics (penicillin/streptomycin) and 1 µg/mL TPCK-treated trypsin]. Afterwards, 50 µL standardized virus in VGM-T was added to each plate or VGM-T to cell control wells. The virus stocks were standardized by FFA to determine Focus Forming Units per milliliter (FFU/mL). Following a 2-hour incubation period at 37**°**C, an 100 µL overlay consisting of equal volumes of 1.2% Avicel RC/CL (45) in 2X MEM containing 1 µg/mL TPCK-treated trypsin, 0.1% BSA, and antibiotics was added to each well. Plates were incubated for 18-22 hours at 37**°**C, 5% CO_2_. The overlays were removed from each well and the monolayer was washed once with PBS to remove any residual Avicel. The plates were fixed with ice-cold 4% (w/v) paraformaldehyde in PBS (10% formalin) for 30 minutes at 4**°**C, followed by a PBS wash and cell permeabilization using 0.5% Triton X-100 in PBS/glycine at room temperature for 20 minutes. Plates were washed three times with PSBT (PBS, 0.1% Tween-20), incubated for 1 hour with a monoclonal antibody pool against influenza A or B nucleoprotein (46) in ELISA buffer (PBS,10% horse serum, 0.1% Tween-80). Following three washes with PBST, the cells were incubated the with goat anti-mouse peroxidase-labelled IgG (Sera Care, Inc., Milford, MA, USA) in ELISA buffer for one hour at room temperature. Plates were washed three times with PBST, and infectious foci (spots) were visualized using TrueBlue substrate containing 0.03% H**_2_**O**_2_**incubated at room temperature for 10-15 minutes. The reaction was stopped by washing five times with deionized water. Plates were dried and the foci enumerated using a CTL Bio Spot Analyser with ImmunoCapture 6.4.87 software (Cellular Technology Ltd., Shaker Heights, OH, USA). The FRA titer was reported as the reciprocal of the highest dilution of serum corresponding to 50% foci reduction compared to the virus control (VC) minus the cell control (CC).

### Antigen-specific Ferret Ig ELISA

Antigen-specific Ig ELISA was performed to assess ferret Ig reactivity against recombinant hemagglutinin (rHA) expressed and purified as previously described (47) and representing the influenza A or influenza B virus strains used for infection. The ELISA was performed using previously described mouse anti-ferret antibodies (5) and similarly to previously described protocols (24). In brief, Immulon 4HBX plates (Thermo Fisher) were coated overnight at 4°C with 1 µg/mL rHA in PBS (50 uL per well) in a humidified chamber and the following day plates were blocked with 200 uL per well of ELISA blocking buffer for at least 1 hour at 37°C. Serum samples were 3-fold serially diluted in blocking buffer in duplicate, and plates incubated for 1 hour at 37°C. Plates were washed four times with PBS, 100 uL per well of mouse anti-ferret IgG “IgG” (mAb 11E3), or mouse anti-ferret IgKappa/IgLambda (mAb 8H9 + mAb 4B10, hereafter referred to as “totIg”) each diluted to 1 µg/mL in blocking buffer, added, and incubated in a humid chamber for 1 hour at 37°C. Plates were washed five times with PBS before adding 100 uL of HRP-conjugated goat anti-mouse IgG1 (Southern Biotech, Birmingham, AL, USA) diluted 1:4000 in blocking buffer and were then incubated overnight at 4°C. Plates were then washed 5 times in PBS before addition of 2,2’-azino-bis(3-ethylbenzothiazoline-6-sulfonic acid (ABTS) substrate (VWR), and incubation at 37°C for 15 minutes. Colorimetric conversion was terminated by addition of 5% SDS (50 uL per well) and absorbance was measured at 414 nm using a BioTek Epoch 2 Microplate Spectrophotometer (Agilent Technologies, Santa Clara, CA, USA). The endpoint absorbance of each sample was determined on a plate-by-plate basis where the absorbances of all the blank wells were averaged and multiplied by three. The GraphPad^®^ Prism 10 “Interpolate a standard curve” function was utilized to generate a standard curve for each individual sample with the absorbances from the serial dilutions followed by plotting the previously calculated endpoint absorbance onto the curve to find the endpoint dilution of each sample. Duplicate ferret samples were run on separate plates in tandem.

### ELISpot assay

ELISpot were performed using anti-ferret antibodies as previously described (48). Briefly, Multi-ScreenHTS HA filter plates (EMD Millipore, Billerica, MA, USA) were coated overnight at 4°C with 25 µg/mL of rHA representing the influenza A [“CA” (H1pdm09) or “KS” (H3N2)] or influenza B [“BR” (B-Victoria lineage) or “PH” (B-Yamagata lineage)] (48) in PBS (50 µL per well). Additional plates were coated with PBS containing 5 µg/mL BSA alone. Plates were washed three times with PBS before blocking with B cell medium, prepared as previously described (48), for at least 1 h at room temperature. Serially diluted ferret PBMC or splenocytes were then incubated in ELISpot plates for 16-18 hours at 37°C, 5% CO_2_. ELISpot plates were then washed three times with PBS + 0.1% Triton X-100 (Sigma-Aldrich), and after an additional five washes with PBS, mouse anti-ferret IgG (mAb 11E3) and mouse anti-ferret totIg (mAb 8H9 + mAb 4B10) diluted each to 1 µg/mL in blocking buffer (PBS containing 2% BSA [Sigma-Aldrich], 1% bovine gelatin [Sigma-Aldrich], and 0.05% Tween 20 [Thermo Fisher Scientific]) was added and plates incubated for 2 h at 37°C. Plates were then washed three times with PBS + 0.1% Triton X-100 (Sigma-Aldrich), and after an additional five washes with PBS, alkaline phosphatase goat anti-mouse IgG1 (Southern Biotech, Birmingham AL, USA) diluted 1:4000 in blocking buffer was added and plated were incubated for 2 hours at 37°C. Plates were washed five times with PBS and then twice with distilled water (dH_2_O) before addition of 5-bromo-4-chloro-3-indolyl-phosphate/NBT (BCIP/NBT) one-step solution (Thermo Fisher Scientific) and incubation at 37°C for approximately 15 minutes. Development of the substrate was terminated by decanting solution and gently washing plates with dH_2_O. Plates were air dried at room temperature before automated counting using the S6 Ultimate M2 Analyzer and the ImmunoSpot 7.0.28.5 software (Cellular Technology Ltd., Shaker Heights, OH, USA).

### Gene expression in ferret cells

Ferret primers generated to pro-inflammatory (*MCP1, IL-1B, IL-6),* TH1 (*CXCL10*), TH2 (*IL-2*), T-regulatory (*TGFB1*), T-effector (*IL-4, IL-12p40, IL-17*), apoptosis (*Granzyme A*), Type-I/II/III interferons (*IFNA, IFNB, IFNG, IFNL3*), interferon responses (*STAT1, STAT2, STAT3, RIG-I, SOCS3, TSLP)* and housekeeping (*GAPDH*) genes were used to measure differences in expression levels in ferret cells. For genes using TaqMan, probes were modified with 6-FAM, fluorescein amidites (FAM) fluorophore on the 5’ end and a non-fluorescent Black Hole Quencher®-1 (BHQ-1) on the 3’ end. Additionally, a locked nucleic acid at adenine <LNA A> was incorporated in all probes in order to increase template binding strength for real-time PCR (49). Primers and probes for all ferret genes were generated from published ferret sequences (26, 50–52). All primer/probe sets used are shown in supplemental Table S1.

Quantitative real time PCR (qRT-PCR) was performed using an ABI 7500 Fast Dx Real-Time PCR instrument (Applied Biosystems, Waltham, MA, USA). PCR reactions were performed in a 5 uL RNA reaction volume using SYBR™GreenER™ qPCR SuperMix (Applied Biosystems) or SuperScript™ III Platinum™ One-Step qRT-PCR Kit for TaqMan reactions (InvivoGen, San Diego, CA, USA). An RT reaction for 30 minutes at 50°C, inactivation for 2 minutes at 95°C, followed by 40 amplification cycles at an annealing temperature of 50°C. Reactions were performed on three ferrets for each virus and timepoint and the values were normalized by subtracting the mean value of the cycle threshold (C_T_) from that of the C_T_ for glyceraldehyde-3-phosphate dehydrogenase (GAPDH) housekeeping gene (ΔC_T_). The relative levels of gene expression for infected cells were determined by subtracting the individual ΔC_T_ values from that of average ΔC_T_ values of pre-infection cells (ΔΔC_T_) and expressing the final quantification values (2^-ΔΔCT^) as relative fold changes. Genes upregulated >10^4^-fold (>10,000) were given a value of 10,000 and genes downregulated <10^-4^-fold (<0.0001) were given a value of 0.0001.

### Cytokine/chemokine levels in ferret sera by multiplex assay

Ferret serum samples were stored at -80°C until evaluated. Samples were thawed at room temperature for 30 minutes prior to testing by Luminex ferret multiplex assay. The Luminex Assay kit [Ampersand Biosciences (www.ampersandbio.com) Lake Clear, NY, USA] contained ferret-specific antibodies and proteins in a microsphere-based assay and consisted of antigen-specific antibodies covalently coupled to magnetic Luminex beads and biotinylated detection antibodies in a capture-sandwich format. All incubations were performed at room temperature in 96-well plates. Thirty uL of standard, controls, or serum samples (1:5 dilution) were added followed by ten uL of multiplexed capture-antibody microspheres and ten uL of blocking buffer to each well. The plates were sealed and incubated for 2 hours on a plate shaker. The plates were washed 3 times using the Bio-Plex Pro Wash Station [Bio-Rad (www.bio-rad.com), Hercules, CA, USA] followed by the addition of forty uL of biotinylated detection antibody and incubated for 1 hour on a plate shaker. After incubation, 20 μL diluted streptavidin-phycoerythrin was added to each well, thoroughly mixed, and incubated for 30 minutes. Following 3 washes, the beads in the plates were resuspended in 100 μL of wash buffer. After shaking on a plate shaker for 5 minutes, the plate was then analyzed on a Luminex 200 Analyzer [Luminex Corporation (www.luminexcorp.com), Austin, TX, USA]. The cytokine/chemokine results were determined by extrapolating the analyte concentration from the measured mean fluorescence intensity (MFI) value using the standard curve. Results were generated and extracted using the RBM Plate Reader and Plate Viewer analysis software, respectively. Data were exported with values represented to two significant figures and analyzed in Microsoft Excel and GraphPad Prism 10 (GraphPad Software, La Jolla, CA, USA).

### Type-III IFN bioassay

Interferon lambda (IFNL) levels in influenza-infected ferret serum was detected using HEK-λ reporter cells (HEK-Blue^TM^ IFN-λ cells: InvivoGen, San Diego, CA, USA) designed to monitor the activation of the JAK/STAT/ISGF3 pathway induced by type III IFNs. The HEK-λ cells were generated by stable transfection of HEK-293 cells with the human IFNLR and IL-10R receptor genes as well as the human signal transducers and activators of STAT2 and IRF9 resulting in a fully active IFNL signaling pathway. Activation of the pathway was detected since these cells harbored the secreted embryonic alkaline phosphatase (SEAP) under the control of the ISG54 promoter, which is activated by IFNL. Stimulation of the IFNL triggered the JAK/STAT/ISGF3 pathway and induced SEAP production which was measured by a colorimetric assay at 650 nm using Quanti-Blue^TM^ solution according to the manufacturer’s instructions (InvivoGen, San Diego, CA, USA).

Twenty uL of sera from uninfected, IAV and IBV infected ferrets were added in triplicate wells of 96-well tissue culture plate. A tissue culture flask containing HEK-λ cell monolayers was gently rinsed once with PBS and cells were dislodged and suspended in HEK medium (DMEM, 10% FBS, antibiotics) to a concentration of 2.8×10^5^ cells/mL. 180 uL of HEK-λ cells were added to each well containing twenty uL of sample and to serially 1/2-log diluted (0.1 – 1000 ng/mL) recombinant ferret IFNL3 (Kingfisher Biotech, St. Paul, MN, USA). The plates were incubated for 20Hr at 37°C, 5%CO_2_. After 20hr, twenty uL of supernatant from the wells were transferred to a new 96-well plate and 180 uL of Quanti-Blue^TM^ substrate was added and incubated for 2Hr at 37°C, 5%CO_2_. Absorbance was read at 655nm. A sigmoidal 4-point standard curve from 0.1 to 1000 ng/mL was generated using recombinant ferret IFNL3 protein [Kingfisher Biotech (www.kingfisherbiotech.com), St. Paul, MN, USA) and unknown samples were extrapolated from the standard curve.

### Statistical analysis

GraphPad Prism 10 was used for all statistical analyses (GraphPad Software, La Jolla, CA, USA). One-way ANOVA was used to determine differences over time between viruses and significance between groups was determined by 2-way ANOVA analysis. Gene expression analysis was performed from three ferrets per timepoint and outliers were identified by the Grubbs’ test (α=0.05) by Extreme Studentized Deviate method for removal of a single outlier if found. Spearman (1-tailed, 95% confidence) correlation method used for comparison of virus replication to gene expression or protein concentration (multiplex or bioassay) as well as antibody levels (HI or FRA) and protein concentration. The Student’s t-test (unpaired) was used to assess the statistical differences in the gene expression levels in respect to the uninfected controls and for comparing influenza A to influenza B viruses and antibody secreting cells by ELISpot. A p-value of <0.05 was considered statistically significant: * p<0.05, ** p<0.01, ***p<0.001, ****p<0.0001.

## SUPPLEMENTAL DATA

**Supplemental Table S1:**
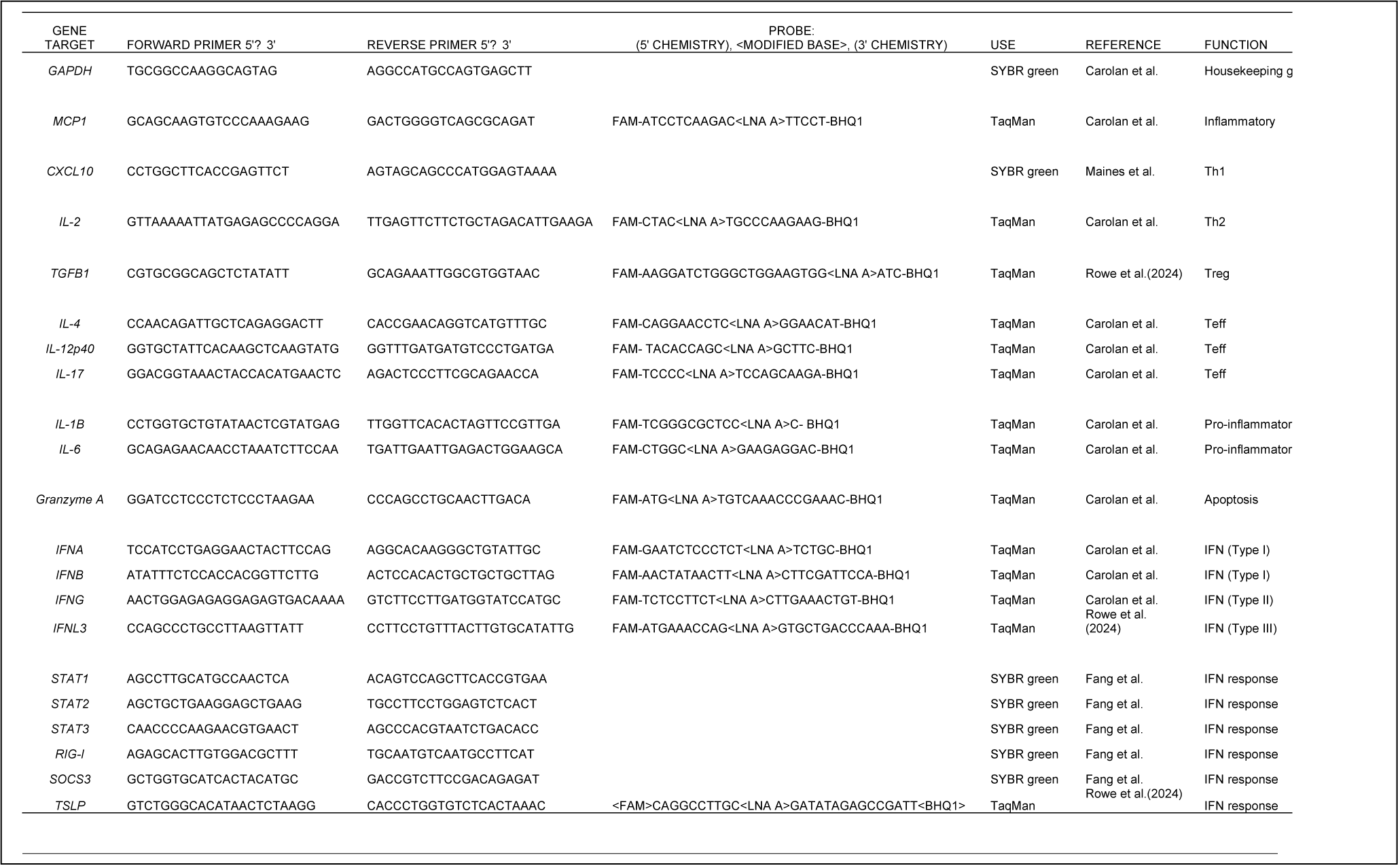
Ferret primers and probes. List of qRT-PCR primers. All primers used in this study are referenced. Forward (5’ to 3’), reverse (5’ to 3’) and probes are listed. Probes using TaqMan enzyme include special chemistry at the 5’-end (FAM) 3’-end (BHQ1) and internally modified bases (LNA A) that were specifically designed for this study to enhance binding and specificity for ferret genes.

**Supplemental Figure S1:**
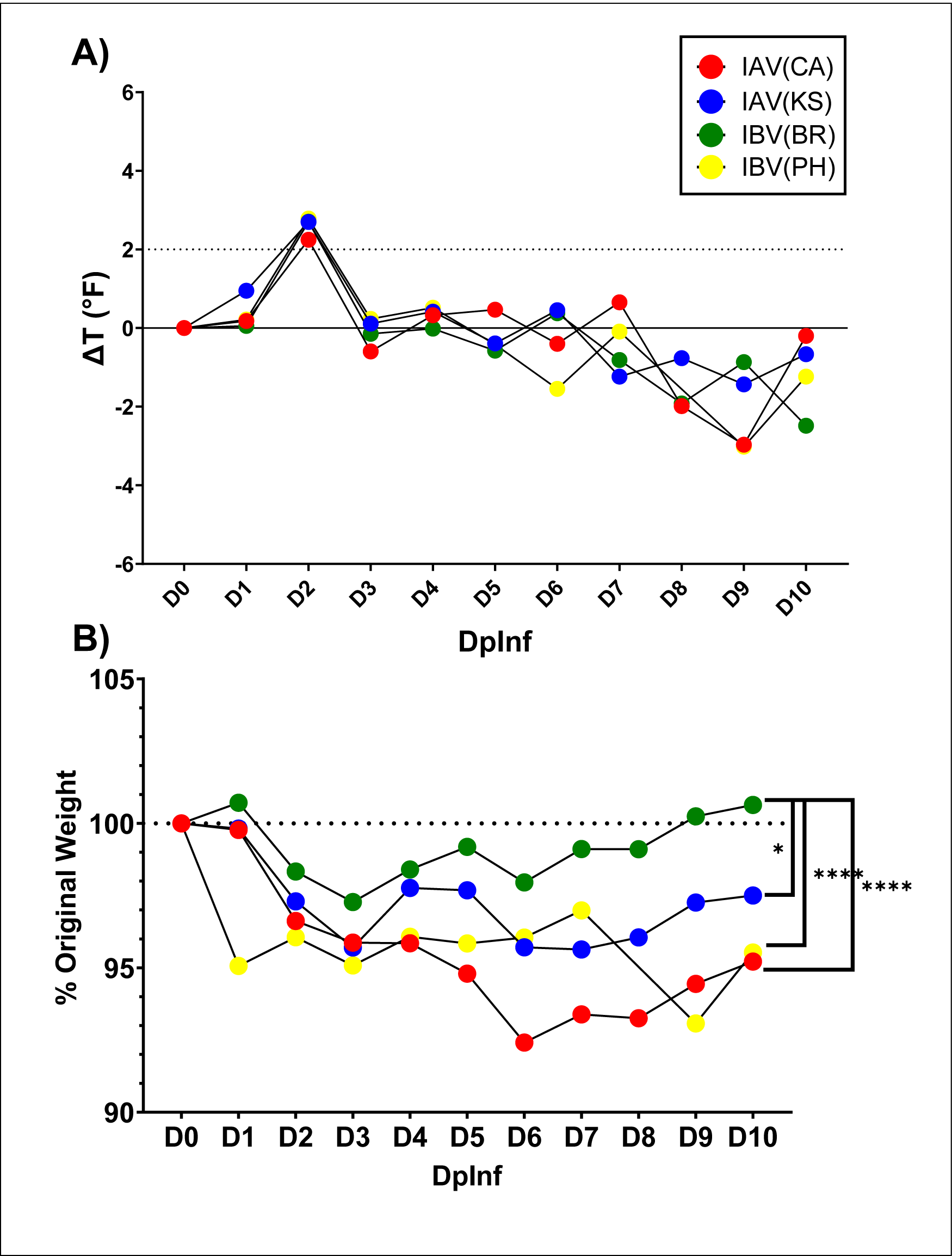
Temperature and weight changes in ferrets following influenza challenge. Clinical parameters in ferrets challenged with IAV and IBV. Red (IAV, H1N1pdm09subtype, CA), blue (IAV, H3N2 subtype, KS), green (IBV, Victoria lineage, BR), yellow (IBV, Yamagata lineage, PH). A) Average body temperature changes from baseline following challenge. Fever indicated by dotted line at +2°F above baseline. B) Weight loss following challenge. Percentage of original, pre-challenge, weight. Significant differences between IAV (CA/KS) and BR as well as PH and BR (2-way ANOVA).

**Supplemental Figure S2:**
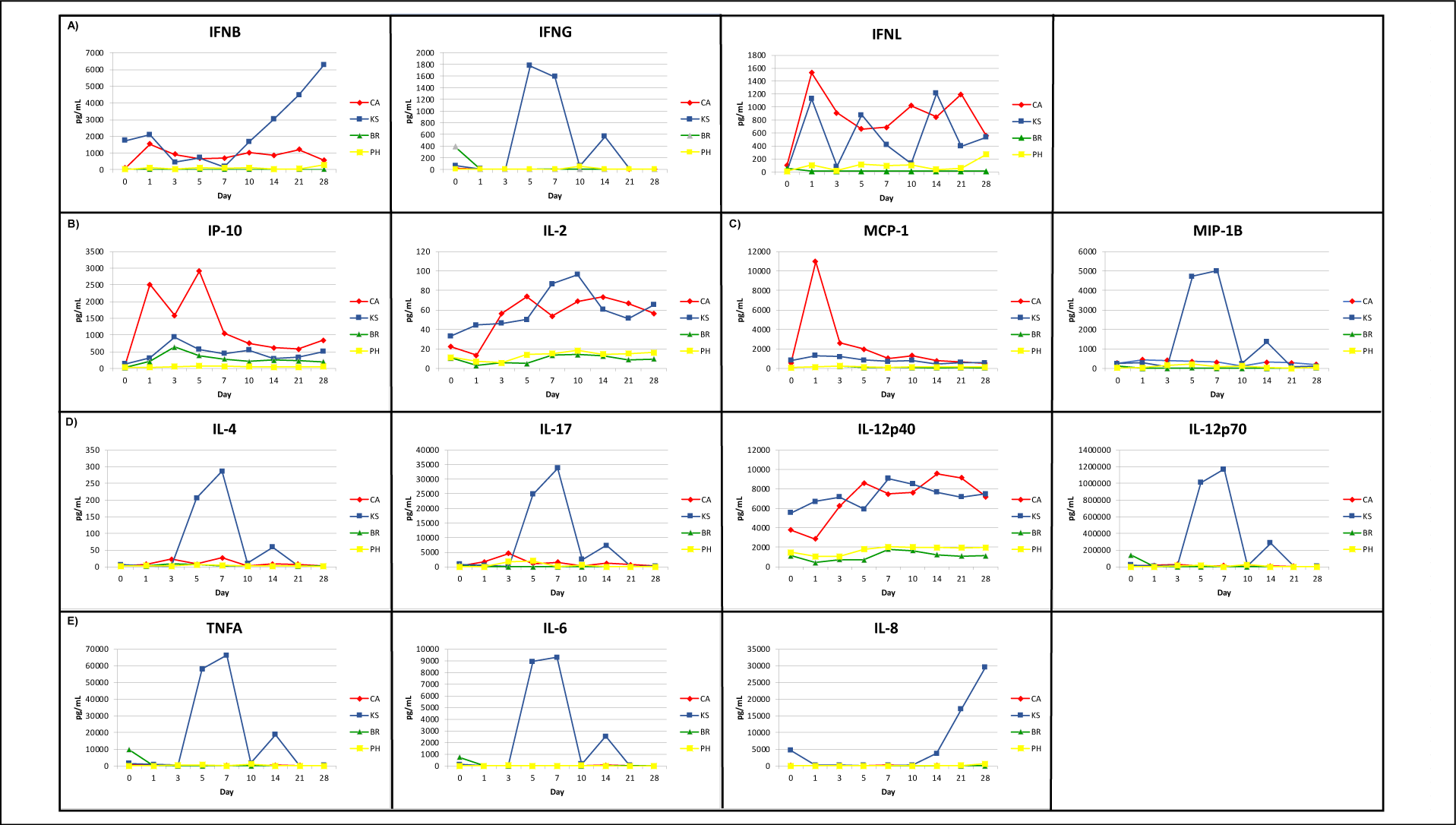
Kinetics of Cytokine and Chemokines in Serum. A) Interferon responses. Type-I (IFNB), Type-II (IFNG) and Type-III* (IFNL). *Bio-assay B) TH1 (IP-10) and TH2 (IL-2) response C)Pro-inflammatory chemokines (MCP-1 and MIP1B) D)T-effector response (IL-4, IL-17, IL-12p40 & IL-12p70) E)Pro-inflammatory cytokine response (TNFA, IL-6, and IL-8).

## AUTHOR CONTRIBUTIONS

Thomas Rowe – designed and conducted experiments, analyzed data. Wrote manuscript

Ashley Fletcher – Multiplexing testing of ferret sera. Data used to help qualify multiplexing assays

Robert A. Richardson – B-cell analysis (PBMC/Splenocytes) – ELISpot & Serum ELISA, manuscript review.

Melissa Lange – serological testing of ferret sera by HI, manuscript review Giuseppe

Sautto – B-cell experiment design and analysis, manuscript review

Greg Kirchenbaum – B-cell experiment design and analysis, manuscript review

Yasuko Hatta – assistance with *In vivo* sample collection and clinical assessments, manuscript review

Gabriela Jasso – assistance with *In vivo* sample collection and clinical assessments.

David E. Wentworth – funding, committee member, and manuscript review

Ted M. Ross – funding, head of committee, experiment, and manuscript review

